# Replay-triggered Brain-wide Activation in Humans

**DOI:** 10.1101/2023.09.14.557724

**Authors:** Qi Huang, Zhibing Xiao, Qianqian Yu, Yuejia Luo, Jiahua Xu, Ray Dolan, Tim Behrens, Yunzhe Liu

## Abstract

The consolidation of discrete experiences into a coherent narrative shape our cognitive map, providing a structured mental representation of our experiences. Neural replay, by fostering crucial hippocampal-cortical dialogue, is thought to be pivotal in this process. However, the brain-wide engagement coinciding with replay bursts remains largely unexplored. In this study, by employing simultaneous EEG-fMRI, we capture both the spatial and temporal dynamics of replay. We find that during mental simulation, the strength of on-task replay, as detected via EEG, correlates with heightened fMRI activity in the hippocampus and medial prefrontal cortex. Intriguingly, increased replay strength also enhances the functional connectivity between the hippocampus and the default mode network, a set of brain regions key to representing cognitive map. Furthermore, during the post-learning resting state, we observed a positive association between increased task-related reactivation, hippocampal activity, and augmented connectivity to the entorhinal cortex. Our findings elucidate the neural mechanism of human replay in both time and space, providing novel insights into dynamics of replay and associated brain-wide activation.

## Introduction

Imagine navigating a new city before bedtime, and then, during sleep, your brain begins to spontaneously recall the route you took throughout the day. This process mirrors the phenomenon known as ‘replay’, a process initially described in rodents, characterized by the fast, sequential reactivation of past experiences, and tied to hippocampal sharp wave ripples (SWRs) ^1,2^. Replay, particularly during rest and sleep, is believed to facilitate hippocampal-cortical dialogue to enhance offline memory consolidation ^3-6^.

Subsequent research has expanded the functions of replay beyond merely facilitating memory consolidation ^7^. Replay also helps organize experiences based on a cognitive map, such as spontaneously representing the rules, or depicting the short-cuts in a maze ^8-10^. Replay has been shown to support reminiscing about past experiences ^9,11,12^, understanding the present ^8,13^, and planning for the future ^14,15^. Replay has been detected not only in the hippocampus but also in other brain regions ^16^, including the visual ^17^ and entorhinal cortex (EC) ^18^, with these replays occurring either in coordination with or independently from hippocampal replay. However, the restricted spatial coverage afforded by invasive neural recording means that the precise pattern of whole-brain activation coinciding with replay events remains largely obscure.

In humans, offline memory reactivations ^19-21^ and recently also signs of sequential replay ^9,22^ have been observed with noninvasive neuroimaging, findings that align with those observed in rodent studies ^23^. During planning, Kurth-Nelson, et al. ^24^ reported fast neural sequences with a time lag of 40 ms using Magnetoencephalography (MEG). The state-to-state time compression seen in these sequences resembled those observed in rodent replay ^4,6^. During rest, reverse replay of past experiences, has been selectively linked to value learning. This was identified using both MEG ^9,14^ and Electroencephalography (EEG) ^25^, a finding consistent with the animal literature ^12^. While M/EEG provides valuable insights into the rapid dynamics of replay, it does not offer precise information about the source of replay signals.

Human functional magnetic resonance imaging (fMRI) has been used to localize sequential neural replay to specific brain regions ^22,26,27^. Wittkuhn and Schuck ^26^, employed fMRI to index the sequence of predictive probabilities within a time repetition (TR), reporting sub-second activations of visual stimuli in the occipital-temporal cortex on-task. During rest, Schuck and Niv ^22^ reported positive correlation between the frequencies of transitions between decoded states and the expected distances between these states in the hippocampus. However, fMRI is limited in its ability to discern the directionality and speed of replay ^26^, characteristics that are likely important given that previous human M/EEG studies ^9,14,28,29^, as well as animal research ^5,12^, have shown a correspondence to different functional aspects of replay. To date, no study has been able to simultaneously record replay events and capture high spatial resolution, whole-brain activity in humans (cf. related work by Higgins, et al. ^30^ on MEG source localization).

Neural replay is implicated in the formation of cognitive map, which not only serve as repositories of experience, but also help rearrange and reconfigure disjointed experiences into a coherent narrative, similar to how our brains reorganize non-chronological movie scenes ^13,31,32^. It is important to understand how replay dynamics relate to wider brain activation, where the default mode network (DMN) in particular, is thought to play a key role. The DMN is a set of brain regions that exhibits increased activity during rest ^33^ or internal cognition ^34^, such as mental simulation or imagination ^35^. It is postulated to encode our knowledge of the world, or cognitive map ^36-38^. However, the dynamics of replay and the DMN in both task and rest remain underexplored because hippocampal replay is temporally transient, while the DMN is spatially distributed. Neither M/EEG nor fMRI is sufficient to capture both neural processes at the same time.

Simultaneous EEG-fMRI recording offers a unique opportunity to address this in greater depth ^39-41^, as the fine temporal resolution of EEG can capture fast neural replay and provide a timestamp for replay events to probe brain-wide activation in fMRI. In this study, we employ simultaneous EEG-fMRI recording during a sequence mental simulation task to probe hippocampal replay and its connectivity to other brain regions, particularly the DMN. The task is designed in a way that subjects build a linear structure (i.e., cognitive map) out of piecemeal experiences. We also include the rest period before (PRE) and after (POST) the task to probe spontaneous task-related reactivation.

## Results

### Task & analysis pipeline

Using concurrent EEG-fMRI recordings, subjects were tasked to mentally connect dots that were separate in experience but could be linked based on a learned relational structure. Previous studies have shown that a similar task, with two sequences (comprised of six pairwise associations), elicits offline reactivations during rest, which can be detected using either MEG ^9^ or EEG ^25^. The current task is modified to include a directional cue to test if replay directionality during mental simulation is subject to explicit instruction. The task is also simplified to contain only one sequence of four objects.

The task starts with a functional localizer session (Fig. 1a), used to train decoders, during which subjects were presented with one of the four images. They were encouraged to think about the image’s semantic content and were later asked to determine whether the following text matched the preceding image. As in previous studies, subjects were unaware of task-related information during the functional localizer session. This session was used to train decoders for both EEG ^25^ and fMRI signals ^26^. After this, three pairwise associations were presented in a randomized order (e.g., 1→2, 3→4, 2→3), and subjects were required to mentally link the associations into a sequence (i.e., 1→2→3→4), a process we term sequence learning. Only those subjects who achieved at least 90% accuracy in the last learning run proceeded to the cued mental simulation task. A resting state period was included both before (PRE Rest) and after learning (POST Rest). After that, subjects were asked to mentally simulate the sequence in either a forward or reverse order, based on the cue (1 →, forward; ← 4, backward). Our subsequent analyses included 33 subjects who completed all task sessions with simultaneous EEG-fMRI recording.

**Fig. 1.**
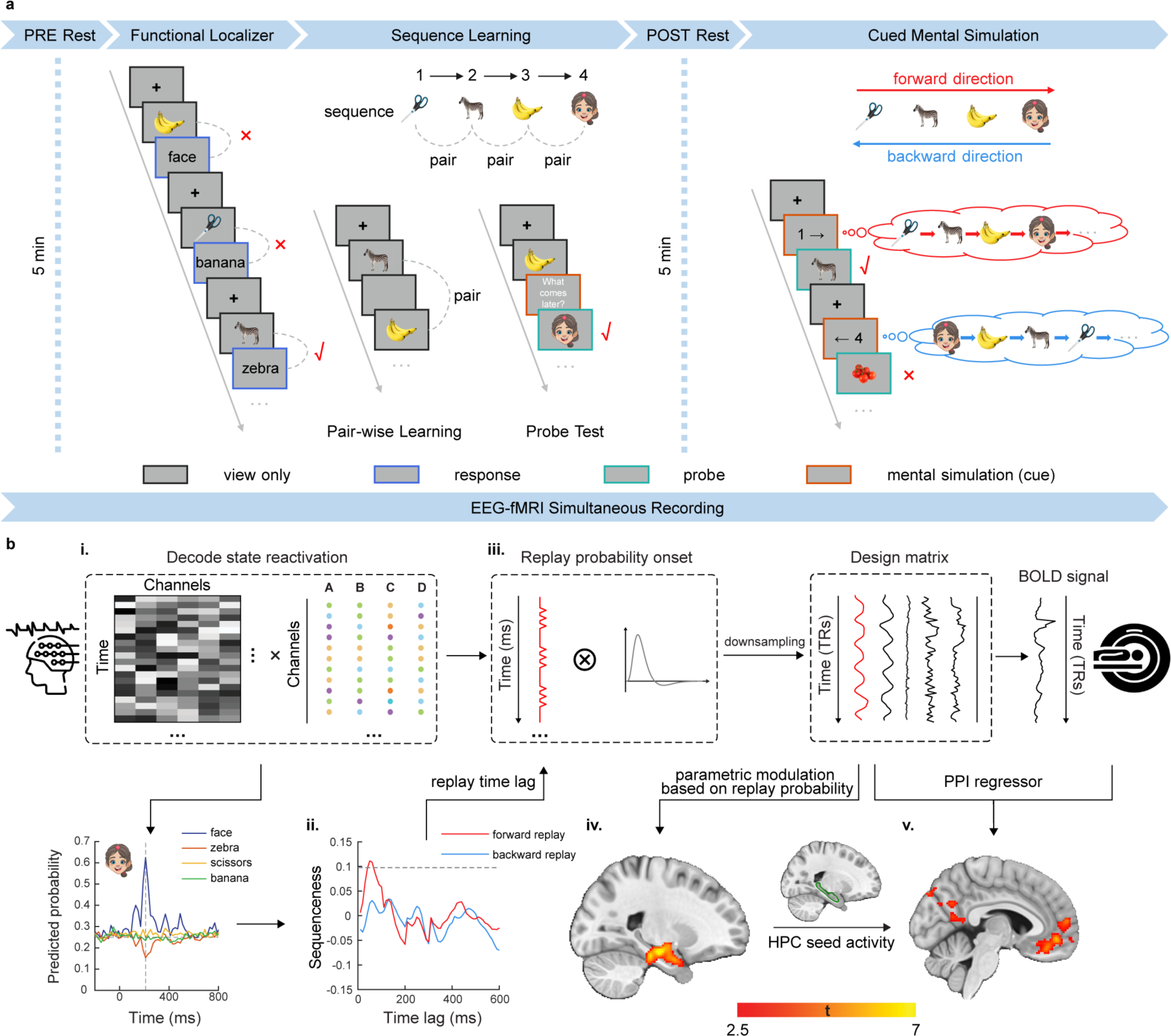
Task schematic and analysis pipeline. **a**, Subjects, undergoing simultaneous EEG-fMRI recordings, were required to construct a sequence by learning pairwise associations of four visual stimuli. They were then cued to mentally simulate the learned sequence in either a forward or reverse order. As in previous replay studies ^9,14,28,30,44,45^, stimuli were first presented in a random order during functional localizer phase, prior to learning. We included a resting state both before (PRE Rest) and after learning (POST Rest) and this allowed us to measure changes in spontaneous neural activity induced by learning. **b,** Details of the simultaneous EEG-fMRI analysis framework for studying replay. **i.** EEG-based stimuli classifiers were trained using whole-brain channel features during the functional localizer and later used to decode stimuli reactivations during specific task phases, such as rest or during mental simulation. **ii.** Temporal Delayed Linear Modelling (TDLM) was applied to the decoded time series to measure the sequential reactivation of states (e.g., visual stimuli) separately for forward and reverse order ^42^. **iii.** After identifying a time lag of interest (e.g., the peak of sequenceness), we derived an EEG-based replay probability time course. This was then convolved with the hemodynamic response function (HRF) and down-sampled to match the fMRI time resolution, serving as an additional regressor in an fMRI-based GLM analysis. **iv.** Based on the new GLM, we determined when (via EEG) and where (via fMRI) replay occurs. **v.** Using an fMRI-derived ROI (green trace, hippocampus), this EEG-based replay probability can be used (by multiplying with ROI neural activity) to detect changes in functional connectivity with other brain regions as a function of replay probability (i.e., psychophysiological interaction, PPI). Data shown here (decoding, EEG replay and coupled fMRI pattern) are from representative subjects. Results are presented with *Punc.* < 0.01 for illustrative purpose and reported using the MNI coordinate system.

Utilizing the fine spatiotemporal resolution offered by simultaneous EEG-fMRI, our goal is to determine when and where neural replay occurs in the brain. This involves indexing fast replay events through EEG and imaging replay-aligned brain-wide activation in fMRI. In brief, our analysis pipeline comprises five steps (Fig. 1b). First, we train neural decoding models for each image based on EEG data from the functional localizer session. These models are then applied to decode their neural reactivations during mental simulation and offline resting time. After decoding, we quantify the strength of sequential reactivations (or replay) in a sequence (e.g., 1→2→3→4), separately for forward and reverse order ^42^. If there is significant evidence for replay, we can calculate when such replay occurs and the strength of this evidence. To model this replay probability in fMRI, we convolve it with a canonical hemodynamic response function (HRF), and down-sample it to match the temporal resolution of the fMRI signal. Replay probability can then be encoded as an additional psychological condition using a general linear model (GLM) in fMRI. In addition to localizing replay, we can model its psychophysiological interaction (PPI) ^43^ to explore how functional connectivity between a region of interest (ROI, e.g., the hippocampus) and other brain regions changes as a function of replay probability. Notably, this analysis pipeline is not restricted to replay; we can investigate the spatiotemporal dynamics of any task reactivations in the same way.

### EEG-based and fMRI-based neural decoding

During the functional localizer, subjects were instructed to press key ‘1’ when a text matched the semantic content of its preceding image (congruent condition), and “2” otherwise (incongruent condition). The mean behavioural accuracy was 94.57 ± 0.70%, where chance level is 50%. Following the analysis step outlined above, we trained four separate one-vs-rest logistic regression classifiers based upon EEG data from correct trials, one for each image. As in previous M/EEG-based replay studies ^9,14,24,25,28,30,42,44,45^, we trained EEG decoding models using all available channels as features at a single time bin (10 ms) and tested performance at all time points from 200 ms prior to the stimulus onset to 800 ms post onset (Fig. 2a). The peak cross-validated decoding accuracy was observed at 210 ms post stimulus onset (46.25 ± 0.95%, with a chance level of 25%). To further examine the sensitivity of the classifiers to each image, we analyzed the time course of predicted probability separately for each image (Fig. 2b). All image classifiers showed above-chance probability in predicting the images they were trained on (dark grey lines) and not for other images (lighter grey lines). Based on these results, the image classifiers were trained at 210 ms post stimulus onset for our subsequent EEG-based replay analysis. Note, similar decoding accuracy and temporal dynamics were observed in a pilot subject who performed under both standalone EEG and simultaneous EEG-fMRI settings, indicating consistent neural dynamics across both settings (Supplementary Fig. 1).

**Fig. 2.**
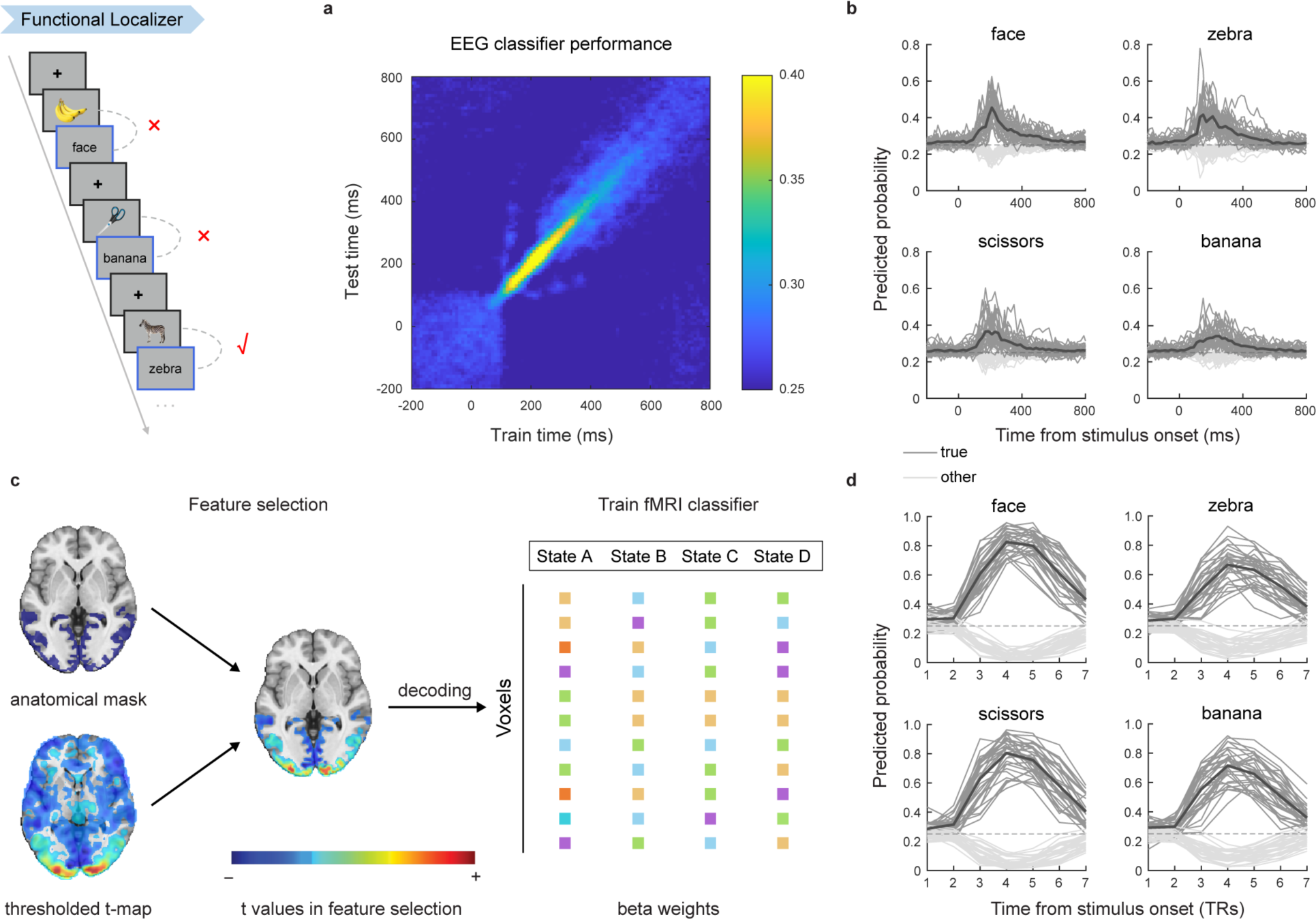
EEG-based and fMRI-based decoding during functional localizer. **a**, The mean cross validated decoding accuracy of EEG-based classifiers. As in previous studies ^9,14,25,28,30,44,45^, classifiers were trained independently at each time point and tested on all time points, starting from 200 ms before stimulus onset to 800 ms post onset (10 ms time bin). Decoding accuracy peaked at 210 ms post-stimulus onset. **b**, The time course (-200 – 800 ms) of mean EEG-based decoding probability trained and tested at the same post-stimulus onset (black line), separately for each stimulus. The dark grey lines represent the decoding probability of a particular classifier for a given image (black line represent the mean probability across subjects), while the light grey lines represent the mean decoding probability of the same classifier for other images. **c**, Feature selection procedure in fMRI-based decoding. Following Wittkuhn and Schuck ^26^, we selected the subject-specific anatomical masks combined with thresholding t-maps (*t* > 3) to identify voxels that selectively response to functional localizer. Note that the result presented here was selected from a representative subject for illustrative purpose only. **d**, The time course (in TR, starting from stimulus onset) of mean fMRI-based decoding probability trained and tested at the same post-stimulus time, separately for each stimulus. The dark grey lines represent the decoding probability of a particular classifier for a given image (black line represent the mean probability across subjects), while light grey lines represent the mean decoding probability of the same classifier for other images.

Contrary to the fine temporal resolution offered by EEG, fMRI is better suited for localizing where in the brain a specific cognitive process unfolds. In fMRI, we found significant activation in the visual cortex when an image was on-screen, with also class-specific activation patterns observed (Supplementary Fig. 2a). Moreover, heightened activation was detected in the temporal cortex and anterior cingulate cortex (ACC) when semantic text was presented (Supplementary Fig. 2b). We have also decoded images based on the fMRI signal and as in Wittkuhn and Schuck ^26^. We first performed feature selection based on anatomical mask and functional *t* map (Fig. 2c), and found a peak cross-validated decoding accuracy of 83.39 ± 1.77%, at the 4^th^ TR post stimulus onset (Fig. 2d), consistent with Wittkuhn and Schuck ^26^.

In both EEG and fMRI-based decoding, the predicted probability for the true stimulus was significantly higher than that of all other stimuli at the same time point (Fig. 2b & d). One advantage of simultaneous EEG-fMRI recording is that it allows us to investigate their activations in response to the same event. We found significant positive correlation across subjects between the decoding accuracy of EEG and that of the fMRI classifiers (robust correlation, *r* = 0.49, *P* = 0.004), suggesting that they are likely capturing corresponding processes.

### Spatiotemporal dynamics of neural replay during mental simulation

Human replays have been found to spontaneously reorganize experience in a manner that corresponds to a learnt relational structure ^9,14,24,25,28,30,42,44,45^. In a sequence learning session, subjects learnt to form a linear sequence consisting of four images based on three pairwise associations experienced in randomized order. During learning, images from pairwise associations were presented serially. Heightened activation in visual cortex, dorsal lateral prefrontal cortex (DLPFC) and hippocampus was evident at the onset of the 1^st^ compared to the 2^nd^ image (Supplementary Fig. 3a). Over the course of learning, hippocampal engagement by the 2^nd^ image increased (*β* = 0.015 ± 0.005, *P* < 0.001, Supplementary Fig. 3b), consistent with its role in associative learning ^37,46^. During the probe phase, a target image was first presented on the screen, then subjects were asked to think of which image comes next. A probe image was then shown, and subjects were required to determine if it was correct or not. We found significant activation in the ACC, DLPFC and insular cortex at the onset of the target image (Supplementary Fig. 3c), and higher ACC activation to the probe image for correct vs. error response trials (Supplementary Fig. 3d). Across all subjects, the mean accuracy of the probe test was 93.86 ± 1.20%, indicating successful learning of the sequence.

After sequence learning, subjects were instructed to mentally simulate the sequence in either forward or reverse order based on a given cue. Subsequently, they were required to identify whether a probe image was part of the sequence. The average behavioral accuracy was 93.59 ± 0.82%. Subjects were also asked to rate the vividness of their subjective experience on a scale from 1 to 4. All participants reported a high level of vividness with a mean rating of 3.35 ± 0.07. No significant difference was found between forward and backward mental simulation in terms of behavioral accuracy (*t*(32) = -0.98, *P* = 0.335) or vividness rating (*t*(32) = 1.50, *P* = 0.143).

At the beginning of each simulation trial, a directional cue, “1→” or “←4” appeared. If replay can be modulated by explicit instruction, we would predict a shift in the direction of replay (if it exists) according to the cue. If replay corresponds to a more unconscious and spontaneous process, then we would expect the direction of replay to be independent of the cue.

In the human neuroimaging literature to date, there are two ways to quantify task-related sequential reactivations or replay in the task. One is Temporal Delayed Linear Modelling (TDLM) ^42^, which calculates the mean sequenceness over all time bins, independently at different speeds (time lags), and separately for forward and reverse direction. This method is mainly used in M/EEG studies 9,14,25,28,30,44,45 but can, in principle, be applied to fMRI data ^42^. The second method uses fMRI data 26, and calculates the regression slope that predicts the position of a state based on the rank of the state probability at each time bin (or TR in fMRI terminology). Fig. 3a, provides an illustration separately for the TDLM method and fMRI-based regression method. Another method for detecting fMRI-based replay is from Schuck and Niv ^22^, calculates the similarity between an hypothesized transition matrix (state distances) and an empirical transition matrix (transition frequency between states) within a brain region of interest (ROI) during rest. In principle, this method can also be applied to the cued mental simulation session.

**Fig. 3.**
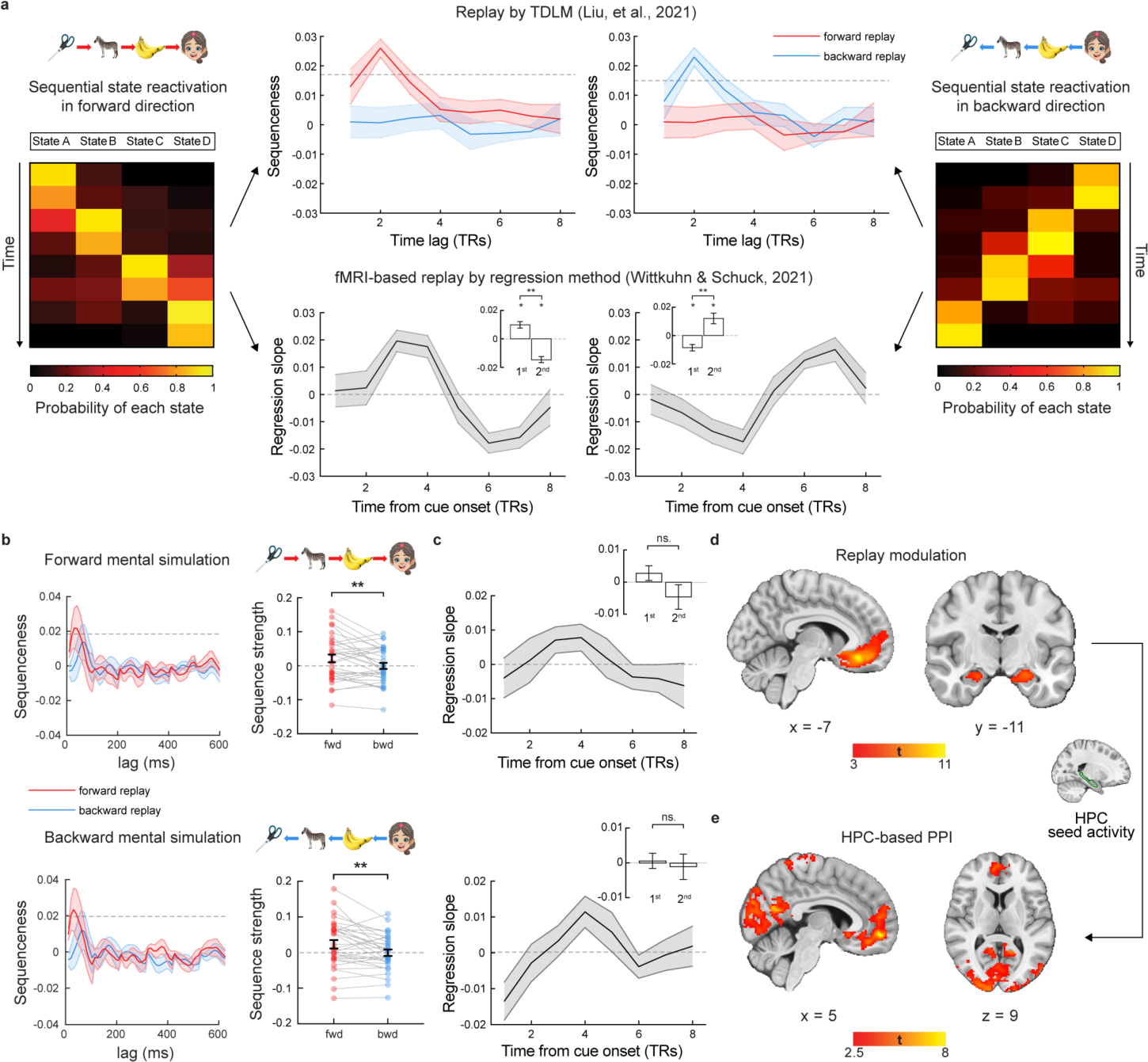
EEG-based and fMRI-based replay during cued mental simulation. **a,** Two analysis methods for detecting replays. One is TDLM ^42^, primarily used with MEG ^9,14,28,30,44,45^, and recently EEG ^25^. The other is a regression method, as per Wittkuhn and Schuck ^26^ and used in fMRI studies ^27^. Note this panel is for illustrative purpose alone. **b**, EEG-based replay with TDLM, separately for forward (cued “1 →”, top row) and backward (cued “← 4”, bottom row) mental simulation conditions. There were significant forward (but not reverse) replays during both forward and backward mental simulation. Sequence strength on the peak time lag (30 ms) is shown on the right, separately for forward and backward mental simulation conditions. The grey dash line represents the premutation threshold, defined as the 95^th^ percentile of the permutated transitions of interest controlling for multiple comparisons. **c**, fMRI-based neural sequence with regression method ^26^, separately for forward and backward mental simulation conditions. There was no significant evidence for sequential activation in the correct order. The bar plot in the upper right corner shows mean slope coefficients for each period. None of these coefficients were significantly different compared to zero. See Supplementary Fig. 4 for assessing fMRI replay using TDLM, as well as results from single subject for illustration purpose. **d**, The parametric modulation of EEG-based replay probability in the whole-brain fMRI during mental simulation showed significant activations in hippocampus and mPFC. We use whole-brain FWE correction at the cluster level (*P* < 0.05) with a cluster-inducing voxel threshold of *Punc.* < 0.001. **e**, The psychophysiological interaction (PPI) between hippocampal activity (anatomically defined) and EEG-based replay probability revealed significant functional connectivity change in mPFC, PCC and visual cortex. We use whole-brain FWE correction at the cluster level (*P* < 0.05) with a cluster-inducing voxel threshold of *Punc.* < 0.01. Each dot is one subject. The grey lines connect results from the same subject. Error bars show SEM. * *P* < 0.05, ** *P* < 0.01, ns., not significant. Abbreviation: HPC - hippocampus.

Using TDLM on the EEG-based decoding, we indeed found selective significant forward replay from 30 to 50 ms time lag in cue “1→” trials (peak at 30 ms lag, *β* = 0.021 ± 0.012, Fig. 3b upper panel), as well as forward replay from 20 to 40 ms time lag in cue “←4” trials (peak at 30 ms lag, *β* = 0.023 ± 0.012, Fig. 3b bottom panel). As the subjects’ task experience increased, their replay strength during mental simulation increased (*t*(32) = 4.18, *P* < 0.001). However, vividness ratings of this simulation, elicited as a subjective measure, were found uncorrelated with replay strength (*t*(31) = -0.55, *P* = 0.585).

In fMRI-based decoding, we also applied TDLM method to the fMRI-based data. While there was a suggestion of replay in some individuals, no significant fMRI-based replays were found across subjects (Supplementary Fig. 4). Likewise, using the regression method ^26^, we did not observe any significant regression slope in either time bin or condition (all *Pcorrected* ≥ 0.06, two-sided one-sample *t*-test against zero, Fig. 3c), nor was there any significant difference between the 1^st^ and 2^nd^ periods (forward: *t*(32) = 1.14, *P* = 0.260; backward: *t*(32) = -0.175, *P* = 0.862, two-sided paired *t*- test, Fig. 3c). Similarly, applying the method from Schuck and Niv ^22^ obtained non-significant fMRI-based replay (Supplementary Fig. 7).

To determine where in the brain on-task neural replay occurs, we identified putative replay events at 30 ms time lag and modelled these in a GLM to predict the fMRI signal. After convolving replay events with the HRF and down-sampling, the replay probability time series was modelled as a parametric modulator of the 10 s mental simulation regressor. We found that the occurrence of replay was associated with activations in both the hippocampus and medial prefrontal cortex (mPFC, Fig. 3d, see also Supplementary Fig. 5a for activations of the mental simulation regressor). This result is consistent with previous findings on MEG replay source localization ^9,14,28,29,44^, suggesting that human replay, as is the case in rodents ^5,47,48^, originates from hippocampus.

We next investigated how functional connectivity between the hippocampus and other brain regions (e.g., DMN) changes in relation to variations in replay probability ^43^. As replay probability increased, there was a significant increase in hippocampal-seed connectivity with DMN, including the mPFC ^49-53^, and the posterior cingulate cortex (PCC) ^52,54^, as well as the visual cortex ^55,56^ (see Fig. 3e and also Supplementary Fig. 5b).

### Spatiotemporal dynamics of learning-induced task reactivation during rest

The findings detailed above indicate that simultaneous EEG-fMRI can index when and where of on-task replay. We next applied this analysis pipeline to rest periods, where, unlike task data, there are no obvious timestamps for specific cognitive processes. Nevertheless, identifying spontaneous replay during rest can provide naturalistic timing information for modelling resting-state activity23. We assumed no significant task-related replay occur during the PRE Rest period, as subjects had not yet experienced the visual stimuli or acquired any structural knowledge. In contrast, during the POST Rest period, after sequence learning, we predicted significant replay ^9^. However, our TDLM analysis did not find evidence of replay in either EEG or fMRI-based decoding during the PRE or POST Rest period (Supplementary Fig. 6). Similarly, using Schuck and Niv ^22^ method to detect replay using fMRI, we found no significant evidence of replay in either the PRE or POST Rest period (see Supplementary Fig. 7).

The relatively simple sequence setup in the current study, which only involved one sequence as opposed to two sequences used in Liu, et al. ^9^, might mean that there is less need for sequential replay during rest ^14,57,58^. Notably, we found that the mean reactivation strength of stimuli, regardless of their sequential order, was significantly higher in the POST Rest compared to the PRE Rest period (*t*(31) = 2.75, *P* = 0.010, two-sided paired *t*-test; Fig. 4a), suggesting enhanced task reactivations following learning.

**Fig. 4.**
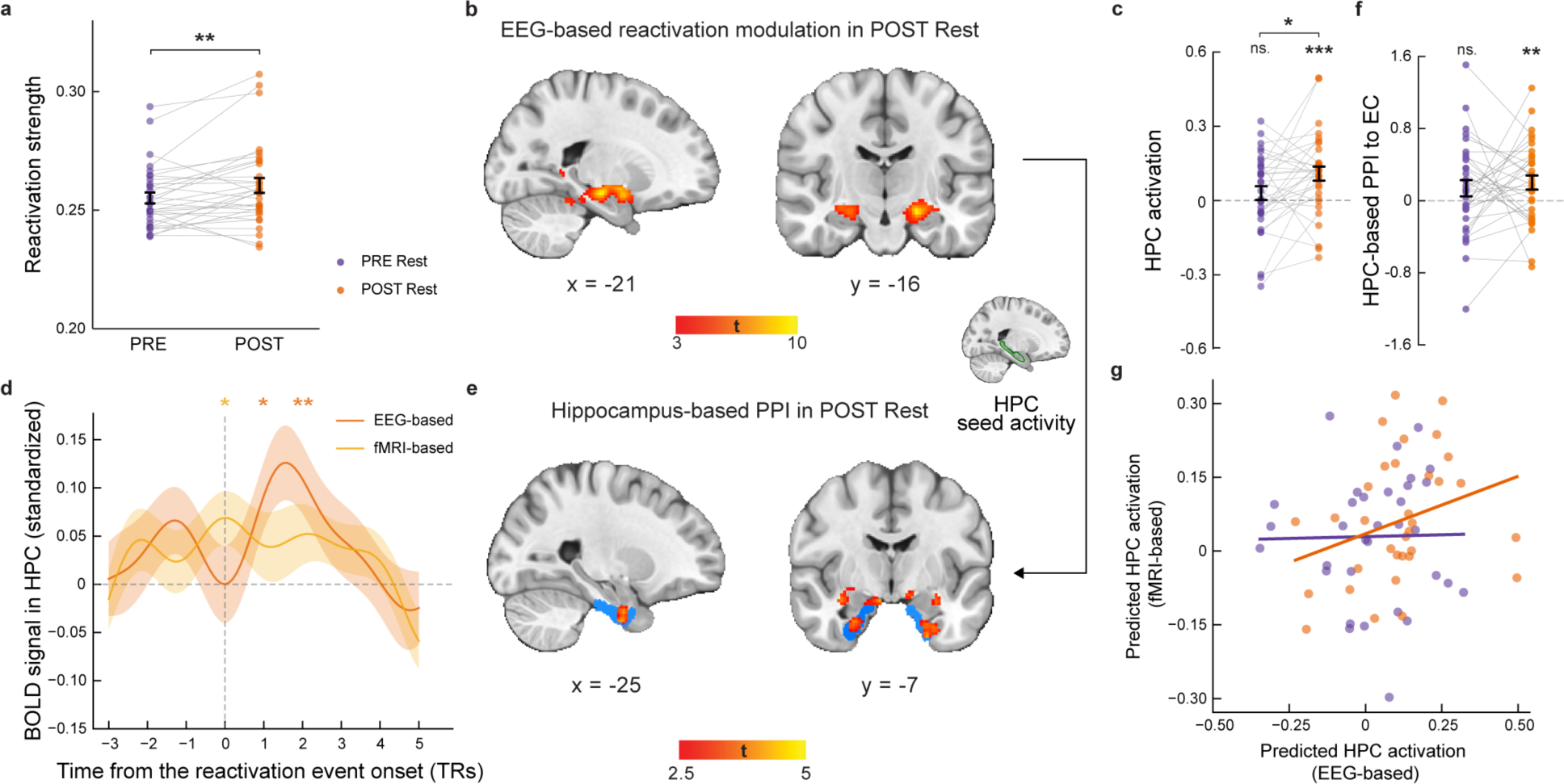
EEG-based reactivation during PRE and POST Rest. **a**, In EEG-based task reactivations, there was a significant higher reactivation strength during POST than PRE Rest. **b**, Parametric modulation of EEG-based reactivation probability in whole-brain fMRI during POST Rest showed significant activations in bilateral anterior hippocampus (whole-brain FWE correction at the cluster level (*P* < 0.05) with a cluster-forming voxel threshold of *Punc.* < 0.001). **c**, ROI analysis. The EEG-based reactivation explained hippocampal activation (anatomically defined) during POST Rest, and it was stronger from PRE to POST Rest in hippocampus. **d**, The task reactivation-aligned BOLD signal in hippocampus during POST Rest. Upon alignment to the onsets of task-related reactivation, we observed a significant increase in hippocampal BOLD activity, peaking at the 2^nd^ TR post-EEG-based reactivation, and also found at onset of fMRI-based reactivation. **e**, The PPI between hippocampal activity (anatomically defined) and EEG-based reactivation probability showed increased functional connectivity with EC during POST Rest. We thresholded at *Punc.* < 0.01, *K* > 10 for visualization. **f**, ROI analysis. The PPI revealed a significant increase in hippocampal-seed connectivity with the EC (anatomically defined) during POST Rest when EEG-based reactivation increased. **g**, There was a positive correlation between EEG-based and fMRI-based reactivation in explaining hippocampal activity during POST Rest, but not PRE Rest. The solid lines reflect the robust linear fit. Each dot is one subject. The grey lines connect results from the same subject. Error bars show SEM. * *P* < 0.05, ** *P* < 0.01, *** *P* < 0.001, ns., not significant. Abbreviation: HPC - hippocampus.

To explore offline reactivation-triggered whole-brain activity patterns, we applied our analysis pipeline to task-related reactivation during rest by summarizing the EEG-based reactivation across stimuli. These reactivation events were then convolved with the HRF, and the ensuing reactivation time series was used as a regressor to explain resting-state fMRI signals. We discovered that higher EEG-based reactivation strength correlated with increased hippocampal activation during POST Rest (Fig. 4b; hippocampal ROI analysis, *t*(32) = 3.83, *P* < 0.001, two-sided one-sample *t*-test), while no activation was identified during PRE Rest at either the whole-brain or the hippocampal ROI level (*t*(32) = 1.08, *P* = 0.287). Moreover, hippocampal activation was significantly stronger during POST Rest compared to PRE Rest (*t*(32) = 2.44, *P* = 0.020, Fig. 4c). As a control analysis, we found that no EEG-based reactivation explained activity in the primary motor cortex (M1), during either PRE or POST Rest or their differences (all *t*(32) ≤ 1.20, *P* ≥ 0.24).

When applying the same analysis pipeline to fMRI-based task reactivation, we found no significant increase in reactivation during POST Rest compared to PRE Rest (*t*(32) = 0.92, *P* = 0.363; Supplementary Fig. 8a). However, in line with EEG-based results, the strength of fMRI-based reactivation was linked to hippocampal activation (ROI level) during POST Rest (Supplementary Fig. 8b-c; hippocampus ROI analysis: *t*(32) = 2.87, *P* = 0.007), but not during PRE Rest (*t*(32) = 1.25, *P* = 0.219). Interestingly, upon alignment to the onset of EEG-based reactivation, a significant increase in hippocampal BOLD activity was observed, peaking at the 2^nd^ TR post-reactivation (*t*(32) = 3.02, *P* = 0.005), indicating a higher sensitivity than that of fMRI-based reactivation (Fig. 4d). This finding suggests that EEG provide an effective means for localizing the timing of spontaneous task reactivation during rest.

We have also examined functional connectivity between the hippocampus and other brain regions as a function of task reactivation during rest (Fig. 4e). The PPI analysis revealed a significant increase in hippocampal-seed connectivity with EC during POST Rest when EEG-based reactivation increased (ROI analysis, *t*(32) = 2.75, *P* = 0.010, Fig. 4f), but not during PRE Rest (*t*(32) = 1.48, *P* = 0.148). No significant results were found when the analysis is done based on fMRI-based reactivation (all *t*(32) ≤ 1.41, *P* ≥ 0.17).

The differences between EEG- and fMRI-based task reactivation raise an intriguing question as to their relationship. While EEG and fMRI-based reactivation time series themselves were not correlated, nor there was a systematic temporal relationship between them either during task or rest (Supplementary Fig. 9a-9c), we found a significant positive correlation of predicted hippocampal activity with EEG-based reactivation and that of fMRI during POST Rest (Fig. 4g, robust correlation, *r* = 0.38, *P* = 0.029), not PRE Rest (*r* = 0.09, *P* = 0.602). This suggests offline task reactivation may align EEG and fMRI-based representation in hippocampus.

## Discussion

Simultaneous EEG-fMRI analysis framework provides a powerful method to study spontaneous replay and associated brain-wide activation. We show that this combined pipeline can detect fast replay events, as reflected in EEG signals (with a 30 ms time lag), and localize these to the hippocampus, as revealed by simultaneous fMRI. We found that replay during mental simulation is a spontaneous process, one that operates independent of explicit task instruction. An increase in EEG indexed replay strength was associated with significant fMRI activations in hippocampus and mPFC, and a significant increase in hippocampal-seed connectivity with the DMN and visual cortex. During rest, we observed a marked increase in EEG-based task-related reactivation from pre- to post-learning periods. Here, an increase in reactivation strength was associated with a significant increase in hippocampal activation and increased hippocampal-seed connectivity with the EC. Combined, these set of findings highlight the utility of simultaneous EEG-fMRI in characterizing the spatial and temporal dynamics of neural replay.

During mental simulation, we found 30-ms-time-lag on-task replay in EEG, which aligns with previously reported human replay speed ^9,14,24,25,45^. Here we now show that the strength of this replay was associated with activation of hippocampus and mPFC, as revealed by fMRI, a finding consistent with previous fMRI replay findings ^22^ and MEG source localization ^9,14,28,29,45^. To rule out any confounding effects due to simultaneous recordings, we conducted the replay analysis after demonstrating that we could reliably decode visual stimuli based on EEG or fMRI signal in the simultaneous EEG-fMRI setting (Fig. 2a, b). The decoding accuracy and dynamics of simultaneous EEG mirrored those observed in the standalone EEG condition (Supplementary Fig. 1), as well as previous M/EEG decoding findings ^9,25^. In fMRI-based decoding, we followed the methods described by Wittkuhn and Schuck ^26^, achieving similar decoding performance. Thus, we find a good alignment with previous fMRI and M/EEG literature on human replay.

An interesting finding in the current study is that the direction of replay is independent of explicit instruction. In previous studies, replay direction was found to change with task demands, such as value learning ^9,14^, probe questions ^29^, and decision-making versus memory preservation ^28^. A common feature of these prior studies is that a shift in replay direction served a specific computational goal. For instance, replay shifted from forward to backward only for a sequence paired with a reward outcome, but not for a neutral sequence ^9^, where reverse replay is hypothesized to solve credit assignment problems ^14,59^. In the current study, verbal instruction alone does not entail any computational demand and the instruction-independent replay pattern suggests that on-task replay may in fact be spontaneous, independent of volition, and conscious mental effort. Consistent with this, we also found no correlation between the subjective rating of vividness of mental simulation and replay strength. While it could be due to ceiling effect of vividness rating, it is also possible that the content of replay is at the semantic level where the vividness of imagery has little impact on replay quality ^60^.

A key advance in our analysis pipeline is its ability to detect brain-wide activation in coordination with replay. During mental simulation, we found that hippocampal connectivity with the DMN increases as a function of replay strength, where DMN is proposed to encode an internal model of the world ^23,32^. When initializing replay, this increase in connectivity suggests a query from the hippocampus to a putative cognitive map, possibly serving to align experiences into an ordered structure. Our results here are consistent with Kaplan, et al. ^61^, who combined electrophysiological recordings of the hippocampus with whole-brain fMRI in anesthetized monkeys, albeit our method has the additional advantage of accessing the *content* of spontaneous task reactivation in humans. During rest, we found increased task reactivation associated with hippocampal activation during the POST compared to the PRE Rest period, with increased hippocampal connectivity to EC linked to increased reactivation probability during POST Rest (Fig. 4d, 4f). The EC is thought to encode task relational structure ^62^, supported by grid cells in animal research ^63^ and grid-like coding in human fMRI studies ^36,37^. We speculate that this increased hippocampal-EC connectivity may underpin memory consolidation during rest, integrating reactivated task information within an established task structure.

Despite the significant increase in task-related reactivation from the PRE Rest to the POST Rest period, we did not find significant sequential reactivation (or replay) of task states during rest. A recent EEG study, employing the same sequence set-up as Liu, et al. ^9^, reported significant reverse replay during rest following value learning, thus demonstrating the sensitivity of EEG in detecting rest-replay ^25^. One plausible explanation for an absence of rest-replay in the current study is the relatively simple sequence set-up, where subjects have less need to replay during rest as the sequence is already well consolidated, as compared to the more demanding task features in Liu, et al. ^9^ and Yu, et al. ^25^. This conjecture is supported by the near-perfect behavioral performance in the final run of sequence learning and is also consistent with Wimmer, et al. ^29^, who reported enhanced mean reactivation, but not sequential replay, for well-encoded memories. This raises an intriguing question regarding the relationship between learning performance and replay strength, particularly considering previous studies found that replay tends to prioritize weakly encoded memories ^57^. It is conceivable that the learning experience and replay strength follow an inverted U-shaped curve, where the strongest replay occurring with intermediate learning experience ^29,64^, a possibility that warrants more detailed investigation.

In the current study, we found higher decoding accuracy for fMRI-based classifiers compared to EEG, possibly due to the much larger feature size. However, the reactivation/replay analysis on the fMRI signal alone was less effective. During mental simulation, following Wittkuhn and Schuck ^26^, we found a qualitatively similar pattern to EEG-based replay, but this was non-significant in the fMRI signal (Figure 3b, right panel). During rest, we found a chance level of decoding accuracy in hippocampus, and non-significant replay in hippocampus and mPFC, when applying the Schuck and Niv ^22^ method (Supplementary Fig. 7). This might reflect that mental imagery is a degraded, fuzzy experience, difficult to detect (Pearson, 2019), and these fMRI-based replay methods ^22,26,42^ are not optimized for discerning replay events in the current study.

A question arising is whether EEG-based and fMRI-based analyses capture overlapping or independent cognitive processes. We found a significant positive correlation between decoding accuracy of EEG and fMRI classifiers during the functional localizer session, suggesting they capture a common process. However, at the level of reactivation dynamics, we found no temporal correlations between EEG and fMRI, neither during mental simulation nor during rest (Supplementary Fig. 9a-c). When we probed the relationship between EEG and fMRI-based reactivation in explaining hippocampal activation, we found a positive correlation during POST but not PRE Rest (Fig. 4g). This suggests that despite the different temporal dynamics between EEG and fMRI activity, spontaneous task-related reactivations align in the hippocampus. Furthermore, a stronger hippocampal BOLD activity when aligning to the onsets of EEG-based, as opposed to fMRI-based, reactivations (Fig. 4d), suggests that EEG may be more sensitive for localizing the timing of spontaneous task reactivations. Together, these findings imply that simultaneous EEG-fMRI can capture spontaneous cognitive processes, even when these are temporally transient or spatially distributed.

Overall, our study experimentally validates an analysis pipeline for studying replay/reactivation alongside whole-brain activation using simultaneous EEG-fMRI. Identifying the putative replay/reactivation events in EEG provides a unique timestamp for imaging brain-wide activation in fMRI. This same analysis pipeline helps bridge between disparate research areas and provides for a comprehensive understanding of the functions of replay in relation to human cognition. This opens exciting new possibilities for future studies, such as investigating hippocampal replay and grid-like coding during cognitive-map-based computation ^9,36^, as well as a richer examination of memory consolidation during sleep ^23^. It enables a more sophisticated understanding of the entorhinal-hippocampal-prefrontal systems underlying inference and generalization.

## Method

### Participants

A total of 40 healthy adults were recruited for the study. All participants had normal or corrected-to-normal vision and no history of psychiatric or neurological disorders. They were screened for magnetic resonance imaging (MRI) eligibility prior to participation. The experiment was approved by the local ethics committee (reference number: PN-202300012), and all subjects provided written informed consent. After excluding subjects with excessive head motion (FD > 0.2) or incomplete participation, 33 subjects were included in the full analysis (age: 22.91 ± 0.33 years, 17 females, 16 males). None of the subjects reported any prior experience with the stimuli or the behavioral task.

### Task

#### Overview of the task design

After completing preparatory work, subjects were taken into the MRI scanner. We began with a short brain localizer, followed by an 8-min anatomical scan and a 5-min resting-state scan, during which subjects were asked to stay awake and focus on a white fixation cross presented on a grey screen. Then, the subjects underwent a series of task sessions: functional localizer, sequence learning, and cued mental simulation. We acquired four functional localizer runs of approximately 12 min each, three sequence learning runs of 6 min each. After sequence learning, we acquired a further 5-min resting-state, again with subjects’ eye open. Finally, we acquired three cued mental simulation runs of about 10 min. The entire experiment lasted, on average, between 2 and 3 hours.

#### Functional localizer

The functional localizer session was designed to train neural decoders on task states. The experiment utilized four visual stimuli (face, scissor, zebra, and banana), which were previously shown to elicit object-specific neural patterns in human brain ^14,44^. Subjects were presented with one of the four images for 1 s and were encouraged to think about its semantic content. Following this, they were presented with text for 1 s, following a 1-2 s blank interval, and asked to determine whether the text matched the preceding image. Subjects were required to press the key ‘1’ if the visual and semantic stimuli were congruent, and ‘2’ if they were not. The key positions were counterbalanced between subjects. Trials were separated by an interval of 1-3 s to allow for sufficient time delay between trials. If participants responded incorrectly, they were given visual feedback for 1 s. Each visual stimulus appeared 72 times with a congruent and incongruent semantic stimulus following. This phase consisted of 288 trials, half congruent and half incongruent. The visual stimuli were presented in a pseudo-random order, with no more than two consecutive presentations of the same stimulus. Subjects with an accuracy rate above 90% would receive a ¥ 20 bonus.

#### Sequence learning

In sequence learning session, subjects were required to build a 4-item sequence (e.g., A→B→C→D) by mentally connecting three pairwise experiences (i.e., A→B, B→C, C→D). The task comprised three runs, each including an associative learning (with three learning pairs) and a probe test. During learning, each trial started with a 300 ms fixation, then stimuli within the learning pair were presented sequentially, one for 1.5 s, with a 1-3 s interval between stimuli. The interval between learning pairs was 5 s. Each learning pair was repeated three times in a run. Subjects were asked to learn associations between stimuli, and their memory performances were probed in the following test. During test, a target stimulus with ‘->…->…?’ cue was presented for 4 s, and subjects were asked to imagine all images that followed the target image. Then after a 1- 3 s interval with a blank screen, subjects were presented with a probe stimulus for 2 s. Subjects pressed key ‘1’ if the probe stimulus followed the target stimulus in the sequence, and ‘2’ otherwise. Key positions were counterbalanced between subjects, and no feedback was given during probe trials. There were 12 probe trials per learning run. The mapping between stimuli (face, scissor, zebra, and banana) and states (A, B, C, D) was fixed within subject but randomized across subjects. Subjects were allowed to proceed if they achieved at least 90% accuracy on the last learning run.

#### Cued mental simulation

Subjects were directed to mentally simulate the image sequence for 10 s, in either a forward (1 →) or a reverse direction (← 4) based upon a directional cue. Then, following a 1-3 s inter-stimulus interval, a probe image was displayed for 2 s. Subjects were required to determine whether the probe image was within the learnt sequence or not. To promote attentive processing, we created four *lure* probe images of the same content with the original ones, but with subtle difference (e.g., orientation of the zebra head, colour of the banana, etc), as well as four new images with different content. The probe images were randomly displayed, with half necessitating a key ‘1’ response if they were in the sequence, and the remaining half requiring a key ‘2’ response if they were not. Key positions ‘1’ and ‘2’ were counterbalanced between subjects. No feedback was provided during probe trials to prevent additional learning. We found no difference of performance in differentiating original and lure images (*t*(32) = 0.62, *P* = 0.541), suggesting the subjects were attentive. After probe test, participants were asked to assess the vividness of their recently performed mental simulation. The task comprised of 96 trials, equally split between forward and backward conditions.

### EEG data acquisition

EEG was recorded simultaneously with fMRI data using an MR-compatible EEG amplifier system (BrainAmps MR-Plus, Brain Products, Germany), along with a specialized electrode cap (BrainCap). The recording was done using 64 channels using the international 10/20 system, with the reference channel positioned at FCz. A drop-down rear electrode was utilized to record electrocardiographic (ECG) activity. EEG data was recorded at a sample rate of 1000 Hz, with the impedance of all channels was kept below 10 kΩ throughout the experiment. To synchronize the EEG and fMRI recordings, the BrainVision recording software (BrainProducts, Germany) was utilized to capture triggers from both the MRI scanner and a stimulus presentation software developed using PsychoPy ^65^.

### MRI data acquisition

All MRI data were acquired using a 64-channel head coil on a research-dedicated 3-Tesla Siemens Magnetom Prisma MRI scanner. For the functional scans, whole-brain images were acquired using a segmented k-space and steady-state T2*-weighted multi-band (MB) echo-planar imaging (EPI) single-echo gradient sequence that is sensitive to the BOLD contrast. This measures local magnetic changes caused by changes in blood oxygenation that accompany neural activity (sequence specification: 46 slices in interleaved ascending order; anterior-to-posterior (A–P) phase-encoding direction; TR = 1300 ms; echo time (TE) = 24 ms; voxel size = 3 × 3 × 3 mm; matrix = 64 × 64; field of view (FOV) = 192 × 192 mm^2^; flip angle (FA) = 67°; distance factor = 0%; MB acceleration factor 2). Slices were tilted for each subject by 30° forwards relative to the rostro-caudal axis to improve the quality of fMRI signal from the hippocampus. For each functional run, the task began after acquisition of the first four volumes (i.e., after 5.2 s) to avoid partial saturation effects and allow for scanner equilibrium. We also recorded two functional runs of resting-state fMRI data, one before and one after the functional localizer and sequence learning task runs. Each resting-state run was about 5 min in length, during which 237 functional volumes were acquired. High-resolution T1-weighted (T1w) anatomical Magnetization Prepared Rapid Gradient Echo (MPRAGE) sequences were obtained from each subject to allow registration and brain-surface reconstruction (sequence specification: 192 slices; TR = 2300 ms; TE = 2.26 ms; FA = 8°; inversion time (TI) = 1000 ms; matrix size = 192 × 256; FOV = 192 × 256 mm^2^; voxel size = 1 × 1 × 1 mm).

### EEG data preparation and preprocessing

EEG data collected inside MRI scanner was contaminated by imaging, ballistocardiographic (BCG) and ocular artifacts. We utilized an Average Artefact Subtraction (AAS) ^66^ algorithm provide by the BrainVision Analyzer software (BrainProducts, Germany) to remove imaging artifacts. Following this, several preprocessing steps were undertaken, which involved removing residual physiological artifacts through the use of EEGLAB ^67^ and custom MATLAB scripts. Specifically, we downsampled the EEG data to a frequency of 100Hz and applied 1 Hz high pass and 40 Hz low pass finite impulse response (FIR) filters. Due to poor signal quality, channel AF3 was excluded from further analysis, and the ECG channel was also excluded. To reduce dependence on the reference electrode position, the average of all electrodes was subtracted from each electrode. The data was then segmented into epochs extending from -200 ms before to 800 ms after the onset of functional localizer stimulus, and from -0.2 s before to 10 s after the onset of the mental simulation cue, without baseline subtraction. Epochs with residual MR artifacts were identified and removed from the dataset. Independent Component Analysis (ICA) was then applied to EEG data, and physiological artifacts ICs belonging to eye, muscle and BCG were meticulously labelled and manually removed.

### MRI data preparation and preprocessing

Results in this manuscript come from preprocessing performed using *fMRIPrep* 21.0.2 (Esteban, et al. ^68^; RRID:SCR_016216), which is based on *Nipype* 1.6.1 (Gorgolewski, et al. ^69^; RRID:SCR_002502). Many internal operations of *fMRIPrep* use *Nilearn* 0.8.1 (Abraham, et al. ^70^, RRID:SCR_001362), mostly within the functional processing workflow. For more details of the pipeline, see https://fmriprep.readthedocs.io/en/latest/workflows.html.

#### Conversion of data to the brain imaging data structure standard

To facilitate further analysis and sharing of data, all study data were arranged according to the Brain Imaging Data Structure (BIDS) specification using *dcm2bids* tool, which is freely available from https://unfmontreal.github.io/Dcm2Bids/.

#### Anatomical data preprocessing

One T1-weighted (T1w) image was found within the input BIDS dataset. The T1-weighted (T1w) image was corrected for intensity non-uniformity (INU) with N4BiasFieldCorrection^71^, distributed with ANTs 2.3.3 (RRID:SCR_004757)^72^, and used as T1w-reference throughout the workflow. The T1w-reference was then skull-stripped with a *Nipype* implementation of the antsBrainExtraction.sh workflow (from ANTs), using OASIS30ANTs as target template. Brain tissue segmentation of cerebrospinal fluid (CSF), white-matter (WM) and gray-matter (GM) was performed on the brain-extracted T1w using fast (FSL 6.0.5.1:57b01774, RRID:SCR_002823)^73^. Brain surfaces were reconstructed using recon-all (FreeSurfer 6.0.1, RRID:SCR_001847)^74^, and the brain mask estimated previously was refined with a custom variation of the method to reconcile ANTs-derived and FreeSurfer-derived segmentations of the cortical gray-matter of Mindboggle (RRID:SCR_002438)^75^. Volume-based spatial normalization to one standard space (MNI152NLin2009cAsym) was performed through nonlinear registration with antsRegistration (ANTs 2.3.3), using brain-extracted versions of both T1w reference and the T1w template. The following template was selected for spatial normalization: *ICBM 152 Nonlinear Asymmetrical template version 2009c* [RRID:SCR_008796; TemplateFlow ID: MNI152NLin2009cAsym]^76^.

#### Functional data preprocessing

For each of the 12 BOLD runs found per subject (across all tasks and sessions), the following preprocessing was performed. First, a reference volume and its skull-stripped version were generated using a custom methodology of *fMRIPrep*. Head-motion parameters with respect to the BOLD reference (transformation matrices, and six corresponding rotation and translation parameters) are estimated before any spatiotemporal filtering using mcflirt (FSL 6.0.5.1:57b01774)^77^. BOLD runs were slice-time corrected to 0.612s (0.5 of slice acquisition range 0 s - 1.23 s) using 3dTshift from AFNI (RRID:SCR_005927)^78^. The BOLD time-series (including slice-timing correction when applied) were resampled onto their original, native space by applying the transforms to correct for head-motion. These resampled BOLD time-series will be referred to as *preprocessed BOLD in original space*, or just *preprocessed BOLD*. The BOLD reference was then co-registered to the T1w reference using bbregister (FreeSurfer) which implements boundary-based registration^79^. Co-registration was configured with six degrees of freedom. Several confounding time-series were calculated based on the *preprocessed BOLD*: framewise displacement (FD), DVARS and three region-wise global signals. FD was computed using two formulations following Power (absolute sum of relative motions)^80^ and Jenkinson (relative root mean square displacement between affines)^77^. FD and DVARS are calculated for each functional run, both using their implementations in *Nipype* (following the definitions by Power, et al.)^80^. The three global signals are extracted within the CSF, the WM, and the whole-brain masks. Additionally, a set of physiological regressors were extracted to allow for component-based noise correction (*CompCor*)^81^. Principal components are estimated after high-pass filtering the *preprocessed BOLD* time-series (using a discrete cosine filter with 128s cut-off) for the two *CompCor* variants: temporal (tCompCor) and anatomical (aCompCor). tCompCor components are then calculated from the top 2% variable voxels within the brain mask. For aCompCor, three probabilistic masks (CSF, WM and combined CSF+WM) are generated in anatomical space. The implementation differs from that of Behzadi et al. in that instead of eroding the masks by 2 pixels on BOLD space, the aCompCor masks are subtracted a mask of pixels that likely contain a volume fraction of GM. This mask is obtained by dilating a GM mask extracted from the FreeSurfer’s *aseg* segmentation, and it ensures components are not extracted from voxels containing a minimal fraction of GM. Finally, these masks are resampled into BOLD space and binarized by thresholding at 0.99 (as in the original implementation). Components are also calculated separately within the WM and CSF masks. For each CompCor decomposition, the *k* components with the largest singular values are retained, such that the retained components’ time series are sufficient to explain 50 percent of variance across the nuisance mask (CSF, WM, combined, or temporal). The remaining components are dropped from consideration. The head-motion estimates calculated in the correction step were also placed within the corresponding confounds file. The confound time series derived from head motion estimates and global signals were expanded with the inclusion of temporal derivatives and quadratic terms for each ^82^. Frames that exceeded a threshold of 0.5 mm FD or 1.5 standardised DVARS were annotated as motion outliers.

The BOLD time-series were resampled into standard space, generating a *preprocessed BOLD run in MNI152NLin2009cAsym space*. First, a reference volume and its skull-stripped version were generated using a custom methodology of *fMRIPrep*. The BOLD time-series were resampled onto the following surfaces (FreeSurfer reconstruction nomenclature): *fsnative*, *fsaverage*. All resamplings can be performed with *a single interpolation step* by composing all the pertinent transformations (i.e., head-motion transform matrices, susceptibility distortion correction when available, and co-registrations to anatomical and output spaces). Gridded (volumetric) resamplings were performed using antsApplyTransforms (ANTs), configured with Lanczos interpolation to minimize the smoothing effects of other kernels ^83^. Non-gridded (surface) resamplings were performed using mri_vol2surf (FreeSurfer).

### Multivariate EEG pattern analysis

Lasso-regularized logistic regression models were trained on EEG data elicited by direct presentations of the images. The preprocessed data from 62 channels were used as input features for the model, which was implemented using the lassoglm function in MATLAB. Each model k had a vector of n + 1 coefficients: one slope for each channel and one intercept. To prevent overfitting, we applied L1 regularization with a lambda coefficient of 0.001. To evaluate the performance of the model, 5-fold cross-validation was employed. The data were randomly divided into five equal-sized subsets, and the model was trained on four subsets and tested on the remaining subset. This process was repeated five times, with each subset serving as the test set once. Decoding accuracy was calculated as the number of correctly classified images divided by the total number of images. We performed decoding at one subject and one time point at a time, repeating the process several times to obtain decoding accuracy for all subjects throughout the entire epoch. We then selected the models corresponding to the time point with the highest accuracy to decode replay or mental simulation.

### Multivariate fMRI pattern analysis

All fMRI pattern classification analyses were conducted using open-source packages from the Python (v.3.9.13) modules Nilearn (v.0.10.0)^70^ and scikit-learn (version 1.1.2)^84^. All multiple comparison correction in fMRI analysis were performed using FMRIB Software Library (FSL, https://fsl.fmrib.ox.ac.uk/fsl/fslwiki/)^85^.

*Feature selection.* Follow Wittkuhn and Schuck ^26^, we combined a functional ROI approach using thresholded t-maps and anatomical masks to identify image-responsive voxels located within a specific brain region. We ran four first-level general linear models (GLMs) for each subject, with one for each of the four cross-validation folds to identify voxels that showed significant activation in response to functional localizer by thresholding t-maps. A first-level GLM was fitted to the training set data (e.g., data from run 2 to 4) of each cross-validation fold and modelled the visual stimulus onset of all corrected trials of functional localizer (1 s for all events). We included wrong trials as a regressor of no interest. All the parameters of GLM analysis were consistent with those utilized in other GLMs (see detail in *GLM analysis* part). These anatomical masks were created based on automated anatomical labelling for brain-surface reconstructions of individual T1w- reference images using *Freesurfer* ^74,86,87^, including the cuneus, lateral occipital sulcus, superior parietal lobule, pericalcarine gyrus, lingual gyrus, inferior parietal lobule, fusiform gyrus, inferior temporal gyrus, the middle temporal gyrus (cf.^29,88^), as well as hippocampus, entorhinal cortex, and para hippocampal gyrus. Only gray-matter voxels were included in the masks^89^. Voxels with t-values above or below a threshold of t = 3 in the anatomical mask for the left-out run (e.g., run 1) of the classification analysis were selected and set to 1 to create the final binarized masks.

*Leave-one-run-out cross validation procedure*. We performed fMRI pattern classification using a leave-one-run-out cross-validation approach, where three task runs (e.g., run 2 to 4) were used for training and the left-out runs (e.g., run 1) used for testing. We trained and tested the classifiers on data obtained from the trials where subjects responded correctly. Four independent one-vs-rest logistic regression classifiers were trained, one for each of the four stimulus classes (*face*, *scissor*, *zebra*, *banana*) and relabeled all other classes to a common *other* category. This process was repeated four times to ensure each task run served as the test set once. All the identical parameter settings were the consistent with those set by Wittkuhn and Schuck ^26^.

To identify the reactivation probability during mental simulation and rest period, we used all the data from functional localizer runs to train the classifiers. We created a new binarized mask for each subject by taking the intersection of the four binarized masks used for cross-validation. The classifiers with the same identical parameter settings as above were trained on the fMRI data. The classifiers were applied to the data during mental simulation (8 volumes per trial), and to data from resting sessions (230 volumes per rest session).

### Temporally Delayed Linear Modelling

We used Temporally Delayed Linear Modelling (TDLM) to measure spontaneous sequential reactivation of four states, either during the mental simulation or rest ^42^. At each time bin during the mental simulation and the two resting sessions, we applied four classifiers to the EEG data and another four classifiers to the fMRI data. Each of these modality data sets contained three [time × state] reactivation probability matrices from the mental simulation (three runs of mental simulation) and two reactivation probability matrices from the resting sessions (PRE and POST Rest).

In a first step, we aimed to identify evidence of state-to-state transitions at a given time lag *Δt*, by regressing a time-lagged copy of one state, *x_j_*, onto another, *x_k_*. In other words, the values of all states *x_k_* at time t are used in a single multilinear model to predict the value of the single state *x_j_* at time *t* + *Δt*:

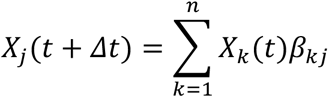

In the second step, we tested for the strength of a particular hypothesized sequence, specified as a transition matrix, T:

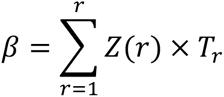

β is the [state × state] empirical transition matrix obtained from previous formula by ordinary least squares regression. *T_r_* is hypothesized transition matrix. In our study, the transition matrices include a forward transition matrix, a backward transition matrix, a diagonal matrix and a constant matrix. Sequenceness, denoted as *Z(r)*, reflected the strength of hypothesized transitions in the empirical matrices, which describe the degree to which representations were reactivated in a task-defined sequential order ^9,14,28,29,44^. *Z_F_* and *Z_B_* represented the forward and backward sequenceness, respectively. By repeating this regression at each time lag, we obtained time courses of sequenceness as a function of time lag. In our research, EEG had a time resolution of 10 ms, the smallest time lag in EEG-based TDLM (Fig. 3b, Supplementary Fig. 6), while fMRI had a time resolution of 1.3s (1 TR), the smallest time lag in fMRI-based TDLM (Fig. 3a, Supplementary Fig. 4 & 6).

We employed a non-parametric permutation-based method to test for statistical significance in this study. For each permutation, sequenceness was averaged across trials within each subject, and then across subjects. The null distribution was constructed by randomly shuffling the identities of the *n* states many times and re-calculating the second-level analysis for each shuffle. We computed a 95% threshold of maximum permutation-derived sequenceness value across all time lags to determine the permutation threshold, which we treated as significant at the level of *α* = 0.05. This approach has been validated in both simulation and empirical data ^9,14,29,42^. As previous human studies have only found evidence for replay with relatively short lags ^9,14,28,42,44^, we visualized results up to a lag of 600 ms in EEG. Simultaneously, we investigated the possibility of detecting replay in fMRI by computing the sequenceness using TDLM with a lag of up to 8 TR.

### Identifying reactivation and replay onsets

To investigate the neural mechanisms underlying replay and task-related reactivation during the mental simulation and rest, we identified the onset of replay and task reactivation for subsequent parametric modulation and psychophysiological interaction analyses (Fig. 1b) ^28,44^. We used classifiers trained on functional localizer to decode the reactivation probability of each visual stimulus, resulting in a [time × state] reactivation matrix. Our analysis revealed a time lag of 30 ms between stimuli that provided the strongest evidence of replay transitions, as determined by TDLM (Fig. 3b). Next, we identified time points during mental simulation and rest, where strong reactivation of one stimulus (e.g., A) was followed 30 ms later by strong reactivation of a structurally-adjacent stimulus (e.g., B). We first generated a matrix *Orig* as

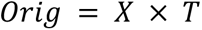

where *X* is the [time × state] reactivation matrix, and *T* is the task transition matrix. The transition matrix *T* defines the mapping between the task state corresponding to column *i* in *X*, and column *i* in *Orig* (specifically, column *i* in *Orig* is the reactivation time course of the state that ‘precedes’ state *i* in *T*). We then shifted each column of *X* by Δ_)_ = 30 ms, to generate another matrix *Proj*,

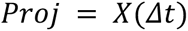

where row *i* of *Proj* correspond to row *i* + 30 ms of *X*. Multiplying *Proj* and *Orig* elementwise, and summing over the columns of the resulting matrix, giving a total of *k* states, to obtain a long [time × 1] vector, *R*. Each element in the *R* indicates the strength of two-state replay at a given moment in time.

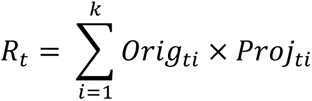

Based on this approach, we calculated forward and backward replay probability onsets for each time point during mental simulation. The replay probability in our study was formed by 30-ms-time-lag forward replay in both forward and backward mental simulation conditions. We convolved the replay probability onsets with the HRF, and downsampled it to the same temporal resolution with fMRI signal. The resulting replay probability onset was an EEG-based replay probability onset regressor. This same method was also employed to the EEG-based task reactivation probability in PRE Rest and POST Rest.

### Detecting neural sequence in task-based fMRI patterns

We employed Wittkuhn and Schuck’s method ^26^ to measure neural sequence in fMRI data during mental simulation. In brief, this involves identifying the relationship between image position in the sequence and task reactivation probability based on an fMRI decoding classifier. Task reactivation probability during cued mental simulation was normalized by dividing them by their trial-wise sum for each stimulus. Subsequently, we conducted a linear regression between the serial position of four images and their normalized decoding probabilities at every TR. The slopes of linear regression were averaged at the subject level for each task condition and each TR. The sign of the mean regression slopes was flipped so that positive values indicate forward ordering and negative values indicate backward ordering. We also performed the two-sided one-sample t-tests to compare the mean regression slope coefficients against zero for each TR and adjusted their *P* values by multiple comparison correction (Fig. 3a for illustration, Fig. 3c for observed data).

It is worth noting that in our study, the [time × state] reactivation probability matrices came from the sequential mental simulation, not sequential visual presentation. Since we could not identify the time at which subjects imagined each image or the speed of imagination, we cannot predict probability differences between two time-shifted events by sinusoidal response functions from Wittkuhn and Schuck ^26^. The junction of the 1^st^ and 2^nd^ period was defined as the point at which the regression slope crossed y = 0 in the forward mental simulation in the positive to negative direction (e.g., junction = 5.2 TR), with the 1^st^ period preceding the junction (e.g., 1^st^ period = [1, 2, 3, 4, 5] TR) and the 2^nd^ period following it (e.g., 2^nd^ period = [6, 7, 8] TR). Slope coefficients were averaged for each task condition and period (Fig. 3a for illustration, the bar plots in the upper right corner of the Fig. 3c for results). We conducted the two-sided one-sample t-tests to compare the mean regression slopes against zero for each task condition and period.

### Detecting sequential replay in rest-based fMRI patterns

We employed Schuck and Niv’s method ^22^ to measure sequential replay in fMRI data acquired during rest. This method involves identifying the relationship between transition frequency between states and state distances. Similar to the aforementioned training of decoding classifiers (refer to *Multivariate fMRI pattern analysis* section), hippocampus and mPFC anatomical masks were created based on automated anatomical labelling for brain-surface reconstructions of individual T1w-reference images using *Freesurfer* ^74,86,87^. We selected the corresponding labels of the bilateral medial orbitofrontal, rostral anterior cingulate, and superior frontal regions for the anatomical mask of mPFC, and the corresponding labels of the bilateral hippocampus regions for the anatomical mask of hippocampus. Considering that these two masks consist of small-quantity voxels, we didn’t employed the thresholded t-map to select image-responsive voxels ^26^. Based on the decoding accuracy, the classifiers trained in mPFC mask at the 5^th^ TR after stimulus onset were chosen to predict the probabilities during mental simulation and rest (see Supplementary Fig. 7).

For each task condition (forward and backward mental simulation) and each rest session (PRE and POST Rest), we selected 230 TRs time series of decoding probabilities. This resulted in 229 state transitions for each condition, allowing us to calculate the transition frequency. Similar to Schuck and Niv ^22^, We conducted a logistic mixed-effects analysis to examine the effects of state distances (hypothesized transition matrix) on transition frequency between states (empirical transition matrix) while simultaneously excluding the different sources of between- and within-participant variability. To compare the models, we employed a likelihood ratio test, comparing a logistic regression model that solely included random effects to a model that also incorporated the state distances regressor.

### GLM analysis

We performed the GLMs to capture the significant event related activations in various sessions: functional localizer (GLM 1), sequence learning (GLM 2, GLM 3), cued mental simulation (GLM 4) and rest periods (GLM 5 for EEG-based reactivation, GLM 6 for fMRI-based reactivation). The fMRI data were smoothed with a 6 mm FWHM kernel before group-level statistics were performed in the GLMs. All images underwent high pass filtering in the temporal domain (width 128s), and autocorrelation of the hemodynamic responses was modelled using an AR (1) model. We included nuisance regressors estimated during preprocessing with fMRIprep: the six rigid-body motion-correction parameters estimated during realignment (three translation and rotation parameters, respectively), *mean White Matter*, and *mean Cerebral Spinal Fluid*. The effect of the experimental conditions on regional blood oxygenation level-dependent responses was estimated with the GLMs. For the group-level analysis, a one-sample t-test was conducted using the whole brain as the volume of interest, and paired t-test was conducted to compare the difference of whole brain activation between PRE Rest and POST Rest. All whole-brain analyses, with the exception for those mentioned otherwise, were thresholded and displayed using a cluster-wise family-wise error (FWE) correction *P* < 0.05, with cluster-forming threshold *Punc.* < 0.001 at the voxel level, as reported by FSL.

#### GLM 1: the activation of images and semantic text in the functional localizer

GLM 1 was employed to find the activation of images and semantic text in the functional localizer session. Each run was modelled with ten regressors, including four regressors to model the onsets of four images, four regressors to model the onsets of four semantic text in correct response trials, one regressor to model the onsets of semantic texts in wrong response trials, and another regressor modelling the onsets of response. To obtain the mean activation of visual processing and semantic processing, we averaged the effect of four images and four semantic texts, respectively (Supplementary Fig. 2a, right panel for images, and Supplementary Fig. 2b, left panel for semantic texts). Furthermore, we identified the specific activation of stimuli by contrasting a specific image or semantic text with the other three images and texts (Supplementary Fig. 2a, left panel for images, and Supplementary Fig. 2b, right panel for semantic texts).

#### GLM 2: the contrast of 1^st^ and 2^nd^ image during sequence learning

GLM 2 was used to examine the differences of activation between the first and second images during sequence learning session. Each run was modelled with two regressors: (1) the onsets of the first image, (2) the onsets of the second image. We contrasted the effect of the first image with that of the second image in the first level GLM (Supplementary Fig. 3a).

#### GLM 3: the activation of target image and probe image in sequence probe test

GLM 3 was designed to investigate the activation of target image and probe image in the sequence probe test. Each run was modelled with four regressors: (1) the onsets of target image (Supplementary Fig. 3c), (2) the onsets of the response, (3) the onsets of correct probe image in all correct response trials, (4) the onsets of wrong probe image in all correct response trials. To access the effect of the wrong probe image in the probe test, we contrasted the effect of probe images between the wrong probe image and correct probe image in the first level GLM (Supplementary Fig. 3d).

### Reactivation and replay onsets modulation analysis

#### GLM 4: the neural correlates of EEG-based replay during mental simulation

To investigate the neural correlates of replay events during mental simulation, we performed a GLM 4 with three regressors. The first regressor represented the onsets of EEG-based replay probability events (See *Identifying reactivation and replay onsets*). We added two more regressors to isolate the unique brain activations associated with replay. The second regressor modelled the duration of mental simulation in all correct-response trials, while the third regressor modelled the duration of mental simulation in all wrong-response trials. These two regressors were modelled as boxcar functions with a duration of 10 s for all trials. We orthogonalized the first two regressors in GLM 4 to remove any shared variances so that the regression coefficients reflected the unique contribution of each regressor in explaining the variances in neural signals.

#### GLM 5: the neural correlates of EEG-based task reactivation in the resting states

To investigate the neural correlates of EEG-based task reactivation during rest, we conducted a GLM 5. We summarized the task-related reactivation probabilities of four states for each time point, resulting in a [time × 1] array of EEG-based reactivation probability onsets. We convolved the reactivation probability onsets with HRF, and downsampled it to the same temporal resolution with fMRI signal. We added it as a psychological regressor to the design matrix of the GLM 5.

#### GLM 6: the neural correlates of fMRI-based task reactivation in the resting states

To investigate the neural correlates of fMRI-based task reactivation during rest, we performed a GLM 6. We summarized the task-related reactivation probabilities of four states for each time point, resulting in a [time × 1] array of fMRI-based reactivation probability onsets. As the fMRI-based reactivation itself has HRF properties, we added it as a psychological regressor to the design matrix of the GLM 6 without HRF convolution.

### ROI analysis

The purpose of ROI analysis in our study is to identify the increased activation during PRE and POST Rest. The beta values at the subject-level for further statistical inference were averaged across all voxels within each ROI. The hippocampus ROI in our study was anatomically defined using a high-resolution probabilistic atlas of Harvard-Oxford Atlas ^90^. The primary motor cortex ROI in our study was anatomically defined using a high-resolution probabilistic atlas of Juelich Histological Atlas ^91^. In the further ROI analysis, any voxels that have any probability of being in the hippocampus and primary motor cortex were included in the ROIs. Two-sided one sample t-tests were performed on beta values for each ROI, rest ression and modality, while two-sided paired t-tests were conducted between rest sessions (PRE versus POST Rest) and modalities (EEG versus fMRI). Additionally, we defined entorhinal cortex ROI by applying 40% threshold to the Juelich Histological Atlas for PPI analysis between hippocampal activity and task reactivation during rest.

### PPI analysis

We performed whole-brain PPI analyses using *nilearn* during mental simulation and rest periods. The first analysis aimed to study replay-triggered brain-wide activation during mental simulation. To achieve this, we used the same hippocampus ROI as the *ROI analysis*. The first PPI model included three regressors for replay onsets: (1) BOLD timeseries extracted from hippocampus (Supplementary. 5b, left panel), (2) the EEG-based replay probability (Supplementary Fig. 5b, right panel), (3) the product of the above two regressors (Fig. 3e).

The second whole-brain PPI analysis aimed to study EEG- and fMRI-based task reactivation aligned brain-wide activation during rest periods. This PPI model included three regressors for task reactivation onsets: (1) BOLD timeseries extracted from hippocampus, (2) the EEG- or fMRI-based task reactivation probability, and (3) the product of the above two regressors (Fig. 4e).

### Cross-correlation

Cross-correlation measures the similarity between two signals as a function of the time lag applied to one of the signals. In the context of EEG-based and fMRI-based decoding probability, during task and rest, we employed the cross-correlation by sliding the EEG time series across the fMRI time series at different time lags and computing the correlation coefficient at each lag. To ensure compatibility between the two signals, we downsampled the EEG time series to the same temporal resolution with fMRI time series before calculating the cross-correlation. We used the cross-correlation function from Liu, et al. ^42^ and the time lag ranging from 1 to 8 TR in our analysis. The peak of the cross-correlation coefficient indicated the point at which the two signals demonstrated the highest degree of similarity. We also performed the two-sided one-sample t-tests to compare the cross-correlation coefficients against zero for each TR and adjusted their *P* values by multiple comparison correction (Supplementary Fig. 9a-c).

### Statistical analysis

Sample sizes were not determined using statistical methods, but were compared with those reported in previous research on replay ^9,14,26,30^. Statistical comparisons were performed using with appropriate inferential methods, as indicated in the figure captions. In cases where multiple hypothesis testing was applicable, we applied the correction method to correct for it ^92^.

## Acknowledgment

Conceptualization, Y.L., Q.H., Z.X., Q.Y., R.D., and T.B.; Investigation, Q.H., Z.X., Q.Y., Y.L., J.X., Y.Q., and Y.Luo.; Writing – Original Draft, Q.H., Z.X., Y.L.; Writing – Review & Editing, Q.Y., Y.L., R.D., and T.B. This study is supported by the National Science and Technology Innovation 2030 Major Program (2022ZD0205500), the National Natural Science Foundation of China (32271093), NSFC (31920103009), the Major Project of National Social Science Foundation (20&ZD153) and Shenzhen-Hong Kong Institute of Brain Science – Shenzhen Fundamental Research Institutions (2022SHIBS0003).

## Data & code availability

The analysis code will be publicly available on https://gitlab.com/liu_lab/EEG-fMRI-replay.git and the data will be publicly available on https://zenodo.org/record/8199536 upon publication.

## Supplementary Figures

**Supplementary Fig. 1.**
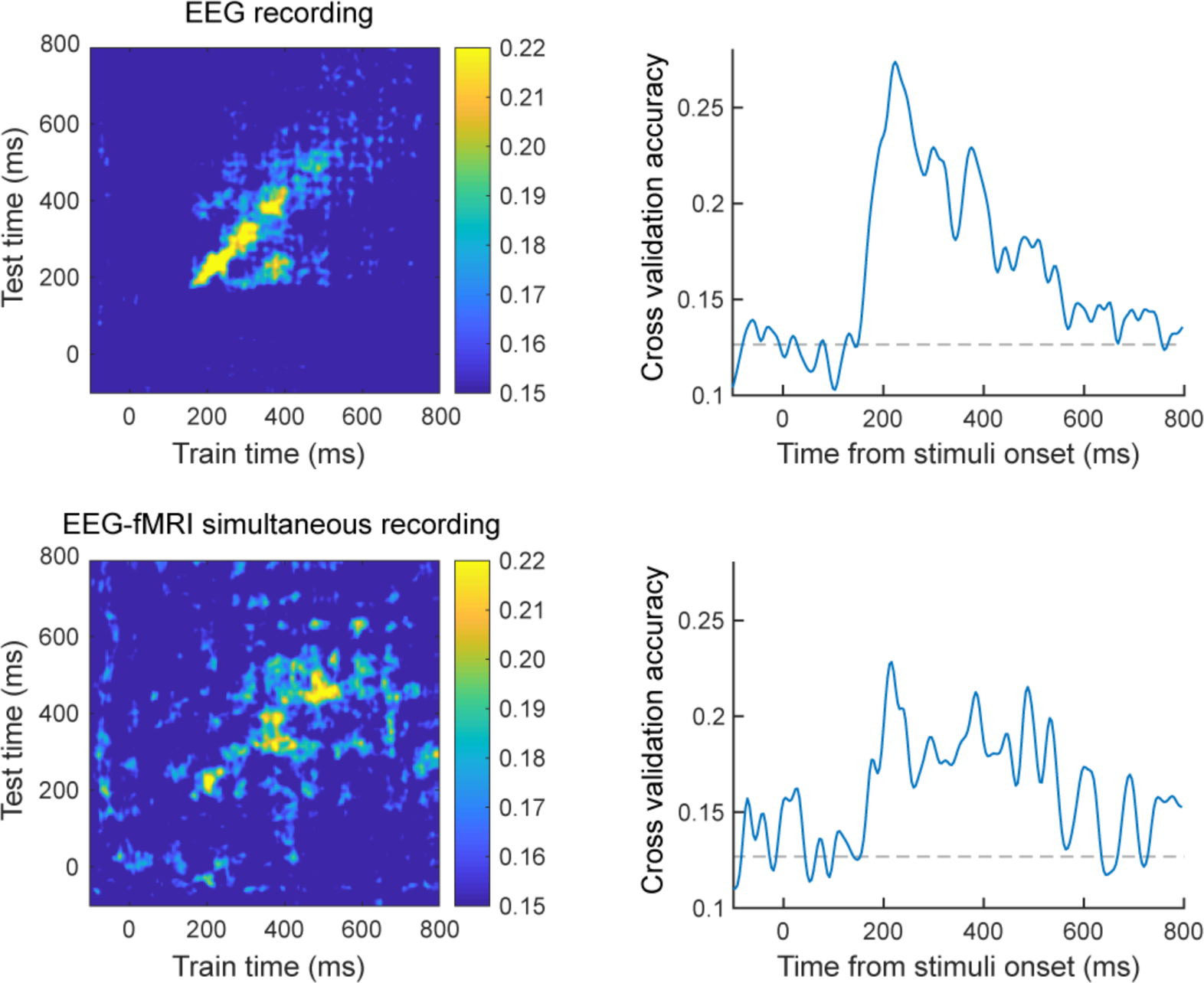
Decoding accuracy between standalone EEG and simultaneous EEG-fMRI from the same subject. The decoding accuracy of EEG signals was compared between the standalone EEG and the simultaneous EEG-fMRI setting from the same subject in a pilot study of functional localizer (with 8 images). Decoding models were constructed separately for each dataset. The figure exhibits the mean decoding accuracy of images for the standalone EEG dataset (upper panel) and simultaneous EEG-fMRI dataset (lower panel), illustrating analogous decoding accuracy and dynamics.

**Supplementary Fig. 2.**
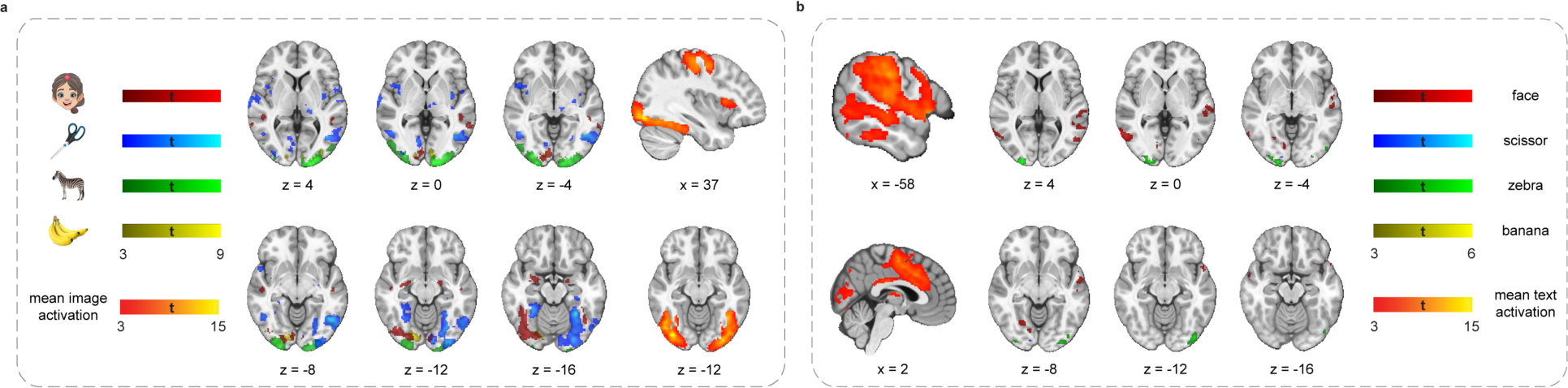
Brain activations of visual and semantic stimuli during functional localizer. **a**, Mean brain activation of all visual stimuli were shown on the right, and localised mostly in visual cortex. The contrast between specific image class and others were shown on the left and revealed class-specific activation pattern (left panel, e.g., face stimuli in fusiform face area). b, Same as panel a, but for the brain activation of text. The mean text activation was shown on the left, localised mostly in temporal cortex, and anterior cingulate cortex (ACC). These results were consistent with previous studies (e.g., ^93^). Compared to the image activation, the contrast of specific text revealed fewer distinctive activation pattern (right panel). We use whole-brain FWE correction at the cluster level (*P <* 0.05) with a cluster-inducing voxel threshold of *Punc*. *&lt*; 0.001.

**Supplementary Fig. 3.**
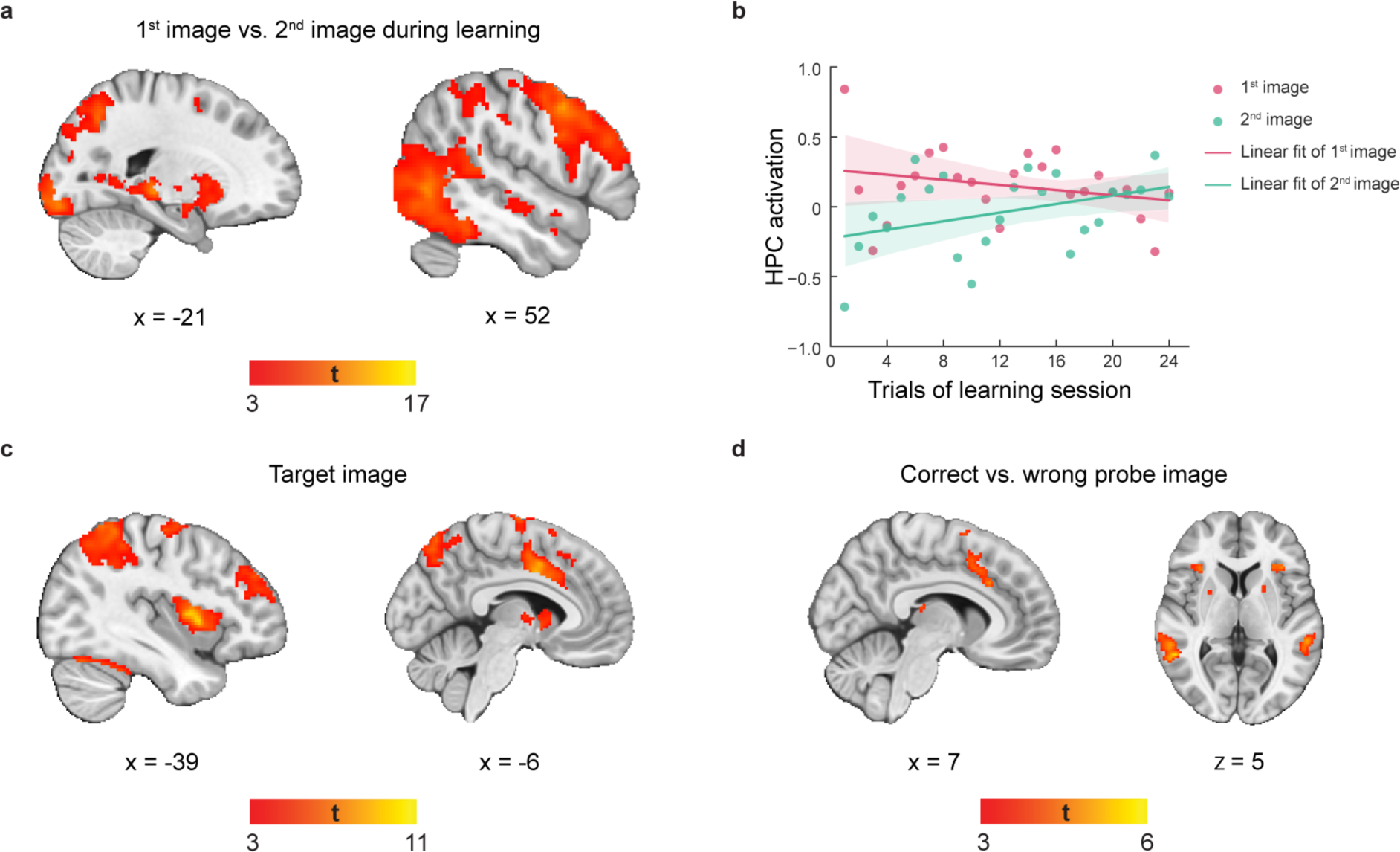
Brain activations during sequence learning and test. During sequence learning, image from pair-wise associations were shown sequentially, and subjects were required to mentally linking three pair-wise associations (e.g., 1→2, 2→3, 3→4, shown in randomized order) to a linear sequence (1→2→3→4). During test session, a target image was shown on the screen and subjects were asked to think about what comes later in the sequence. Then a probe image was shown and subjects needed to indicate whether it was right or wrong. Only trials with correct response were analyzed. **a**, Contrast between 1^st^ and 2^nd^ image presentation reveals activation in temporal lobe (including hippocampus), dorsal lateral prefrontal cortex and visual cortex. **b**, Hippocampal (anatomically defined ROI) involvement as a function of learning trials, separately for 1^st^ and 2^nd^ image. There was a significant increase of hippocampal activation for the 2^nd^ image (linear mixed model, *β* = 0.015 ± 0.005, mean ± SEM, *P* < 0.001), while significant decrease for the 1^st^ one (*β* = -0.009 ± 0.004, mean ± SEM, *P* = 0.039). **c**, During test, there was significant activation in ACC and insular on the presentation of target image. **d**, Contrast between wrong and correct probe image reveals stronger activation in ACC. We use whole-brain FWE correction at the cluster level (*P* < 0.05) with a cluster-inducing voxel threshold of *Punc.* < 0.001. Shaded areas show SEM across subjects.

**Supplementary Fig. 4.**
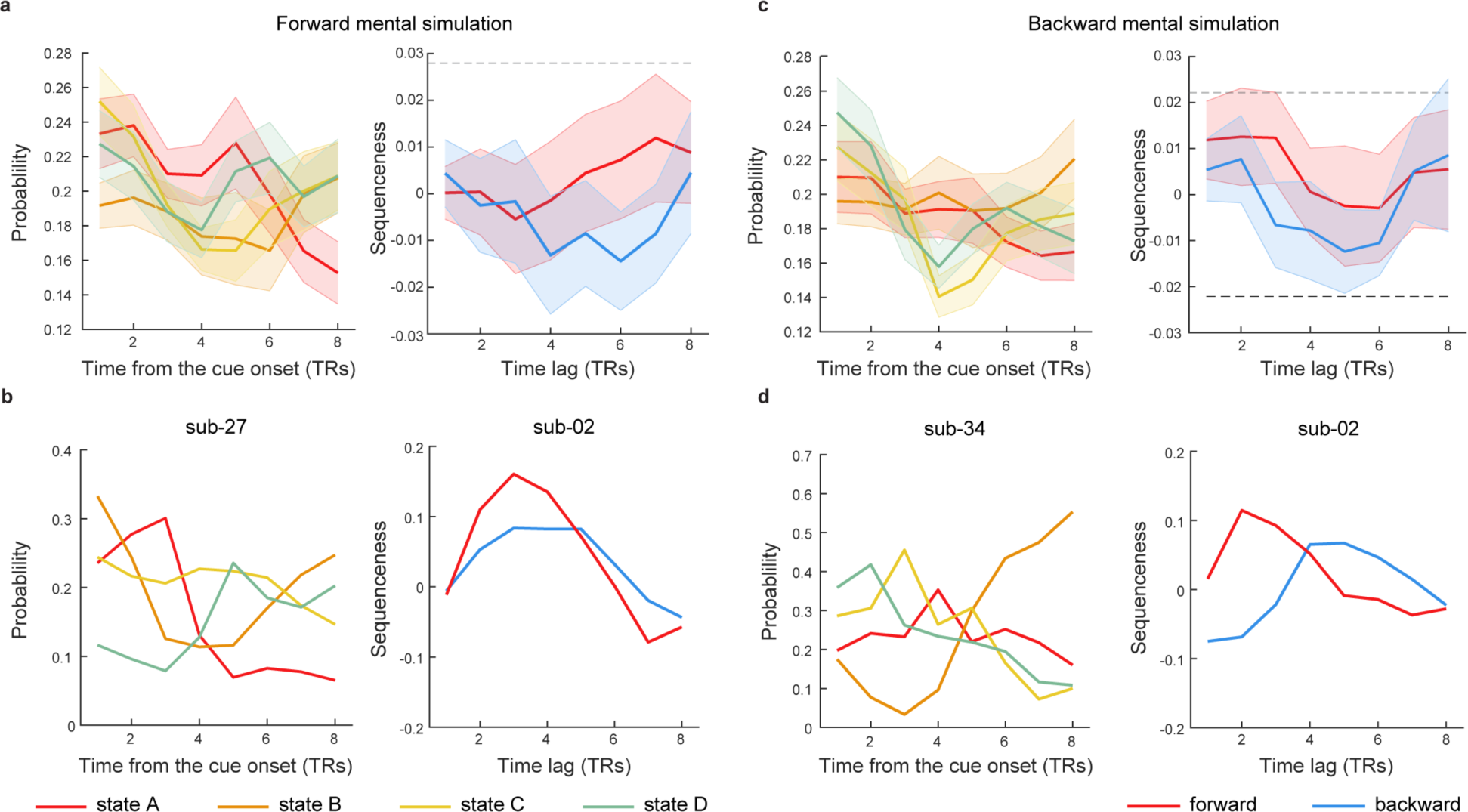
fMRI-based replay during mental simulation. **a**, Mean time course of state reactivation during mental simulation, starting from the onset of cue in forward mental simulation condition (left panel). Based on the state reactivation probability, we detected fMRI-based replay via TDLM. No significant replay was found for forward mental simulation (right panel). **b**, Same with panel **a**, but data from individual subject. **c**, Same with panel **a**, but for backward mental simulation. No significant replay was found for backward mental simulation neither (right panel). **d**, same with panel **c**, but data from individual subject. Shaded areas show SEM across subjects. The grey dash line represents the premutation threshold, defined as the 95^th^ of the permutated transitions of interest controlling for multiple comparisons.

**Supplementary Fig. 5.**
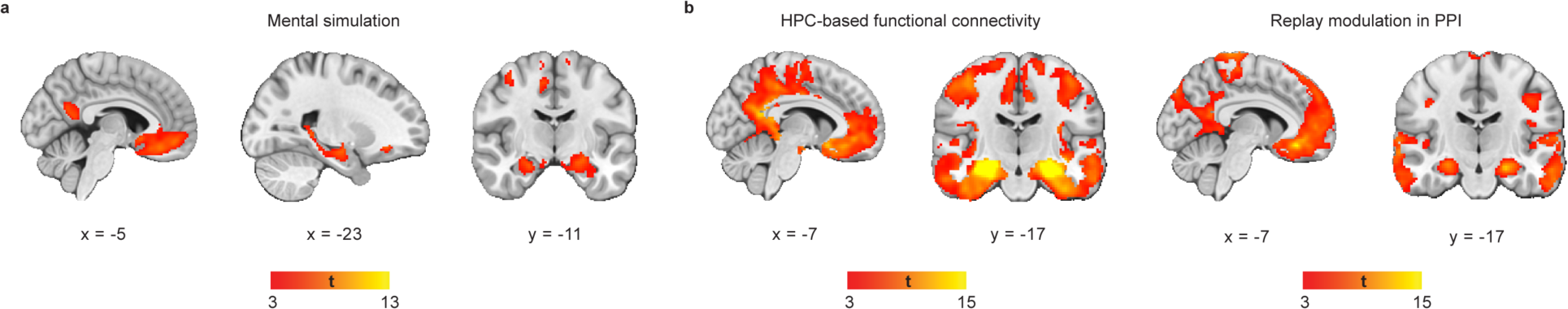
Brain activations and hippocampus-based functional connectivity during mental simulation. **a**, There were significant activations in medial prefrontal cortex (mPFC), posterior cingulate cortex (PCC), orbitofrontal cortex (OFC) and hippocampus during mental simulation. **b**, Main effect of hippocampus (anatomically defined ROI, left panel) and EEG-based replay probability (right panel) in the PPI analysis. Both main effects revealed significant activation in default mode network (DMN), including mPFC, PCC and hippocampus. We use whole-brain FWE correction at the cluster level (*P* < 0.05) with a cluster-inducing voxel threshold of *Punc.* < 0.001. Abbreviation: HPC - hippocampus.

**Supplementary Fig. 6.**
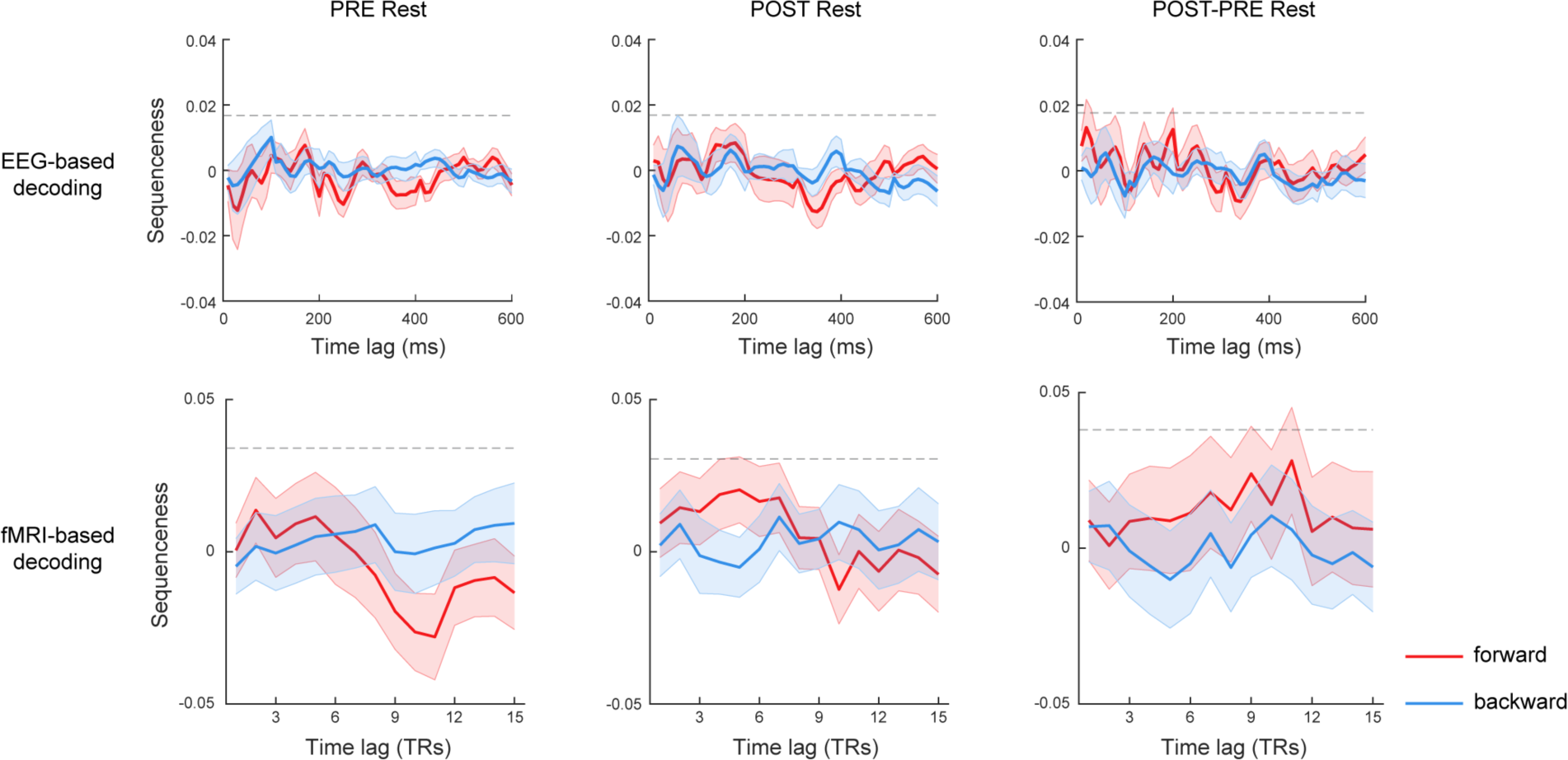
EEG-based and fMRI-based replay during rest. EEG- (upper row) and fMRI-based (bottom row) replay results via TDLM. Results were grouped for different resting time, as well as their contrast (columns). No significant replay was found in any condition. Shaded area show SEM across subjects. The grey dash line represents the premutation threshold, defined as the 95^th^ of the permutated transitions of interest controlling for multiple comparisons.

**Supplementary Fig. 7.**
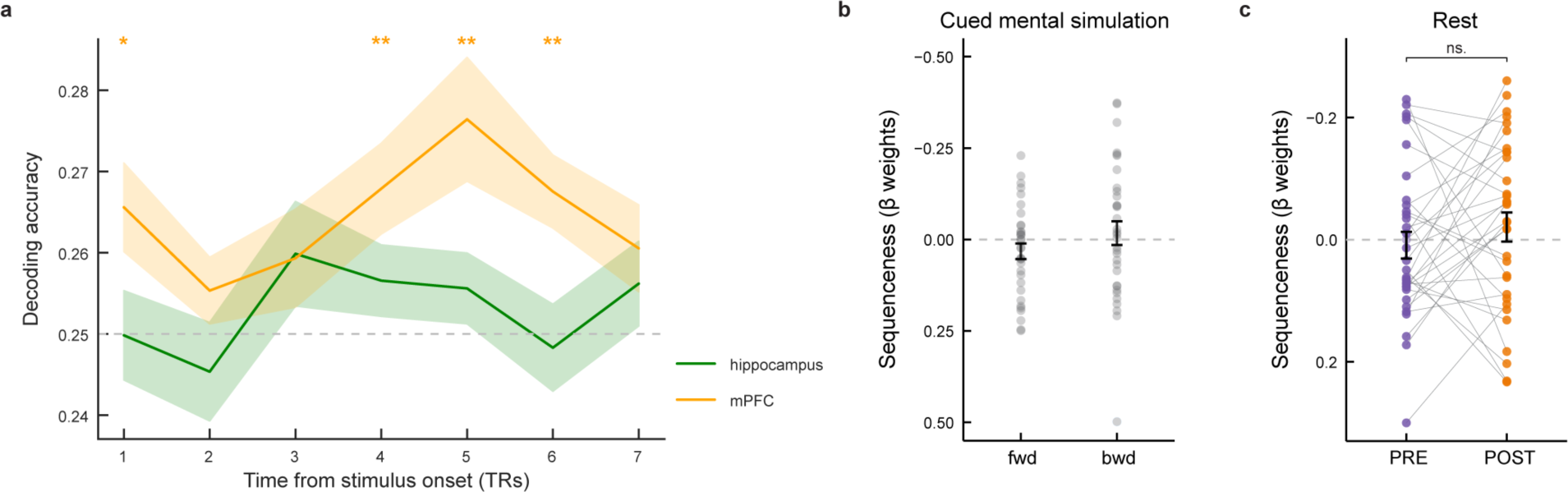
fMRI-based replay during task and rest by Schuck and Niv ^22^ method. **a**, We carried out fMRI-based decoding separately within anatomically defined hippocampus and mPFC masks (see *Detecting sequential replay in rest-based fMRI classifiers part* in the Methods section). In line with previous findings by Wittkuhn and Schuck ^26^, the decoding accuracy within hippocampus did not exceed chance level at any time point from the onset of the stimulus. While, the decoding accuracy within mPFC peaked at the 5th TR following stimulus onset (decoding accuracy = 27.64 ± 0.44%; compared with 25%, *t*(32) = 3.46, *Pcorrected* = 0.005). Shaded areas show SEM across subjects. We then established an fMRI-based mPFC decoding model using data from the 5th TR post stimulus onset in the functional localizer and then employed this model to predict mental simulation (**b**) and two rest sessions (**c**). **b-c**, Employing the methodology developed by Schuck and Niv ^22^, we used a logistic linear mixed model to analyse the impact of state distance on transition frequency during the task (individually for forward and backward mental simulation conditions) and rest. Bars denote the magnitude of fixed effects in the linear mixed model. Error bars depict the SE of the fixed effect estimated at the group level. Each dot represents the beta estimate of state distance for one participant in a logistic regression. Panel **b** showed the replay strength in mental simulation, separately for forward and backward conditions. Panel **c** showed the measured replay in PRE and POST Rest. Model comparisons based on the Akaike Information Criterion (AIC, following Schuck and Niv ^22^) suggested that the integration of the sequenceness regressor did not contribute to a better fit in mental simulation (forward: 2105.52 versus 2107.38; backward: 2051.90 versus 2053.16) or rest session (PRE: 2128.80 versus 2128.38; POST: 2131.05 versus 2132.74). * *P* < 0.05, ** *P* < 0.01, ns., not significant.

**Supplementary Fig. 8.**
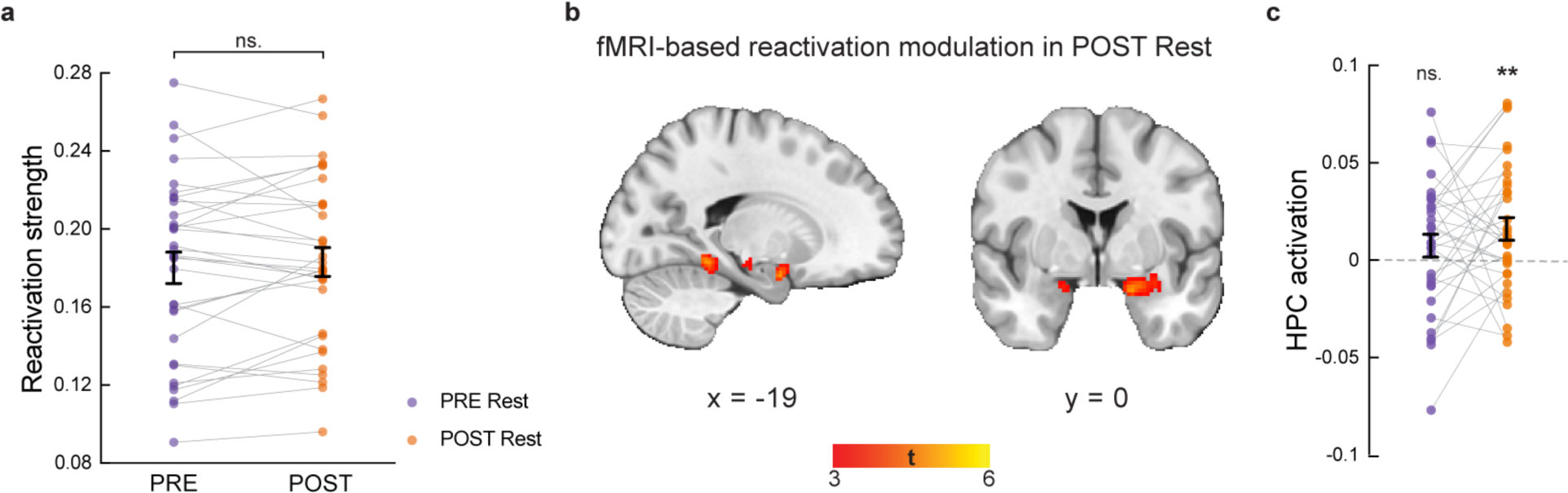
fMRI-based reactivation during PRE and POST Rest. **a**, There was no significant difference of fMRI-based task reactivation between PRE and POST Rest. b, The parametric modulation of fMRI-based reactivation probability during POST Rest. We thresholded at *Punc*. < 0.01, *K* > 10and medial temporal lobe mask for visualization. No cluster survived whole-brain multiple comparison correction. c, ROI analysis. The fMRI-based reactivation explains hippocampal activation (anatomically defined) but not motor cortex (M1, as a control region) during POST Rest. Each dot is data from one subject. The grey lines connect results from the same subject. Error bars show SEM. ** *P* < 0.01, ns., not significant. Abbreviation: HPC - hippocampus.

**Supplementary Fig. 9.**
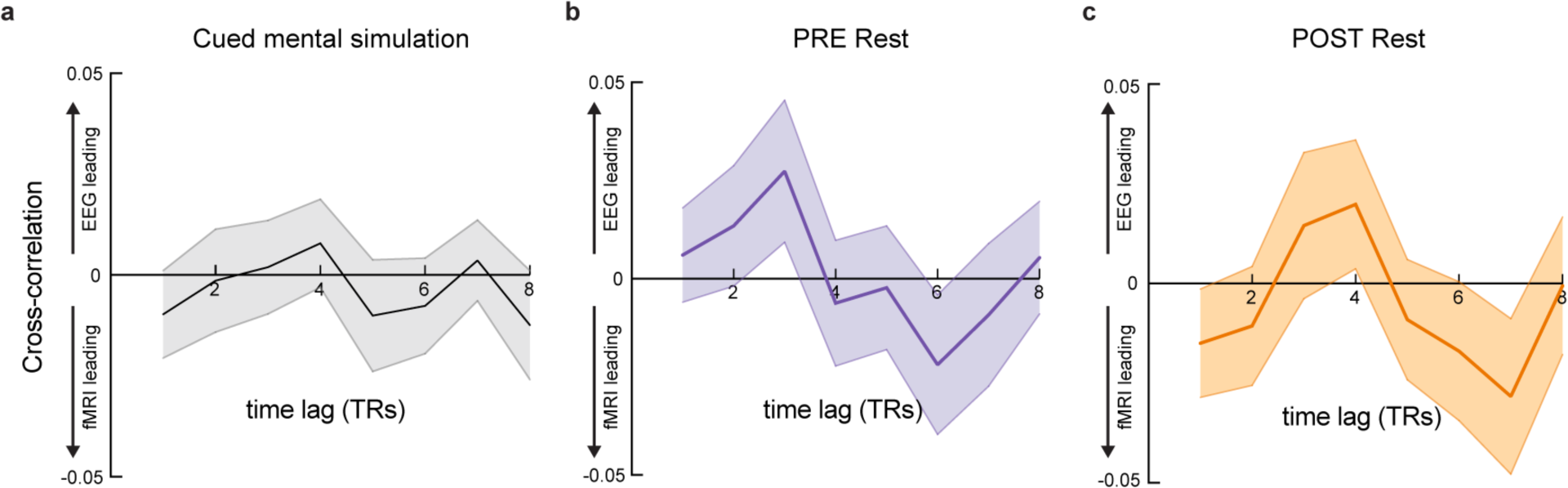
Relationships between EEG-based and fMRI-based reactivation. **a-c**, No temporal correlation was found for EEG-based and fMRI-based reactivation probabilities during mental simulation or rest sessions. Shaded areas show SEM across subjects.

## References

1 Wilson, M. A. & McNaughton, B. L. Reactivation of hippocampal ensemble memories during sleep. Science 265, 676–679 (1994).

2 Nádasdy, Z., Hirase, H., Czurkó, A., Csicsvari, J. & Buzsáki, G. Replay and time compression of recurring spike sequences in the hippocampus. Journal of Neuroscience 19, 9497–9507 (1999).

3 Sutherland, G. R. & McNaughton, B. Memory trace reactivation in hippocampal and neocortical neuronal ensembles. Curr Opin Neurobiol 10, 180–186 (2000).

4 Foster, D. J. & Wilson, M. A. Reverse replay of behavioural sequences in hippocampal place cells during the awake state. Nature 440, 680–683 (2006).

5 Davidson, T. J., Kloosterman, F. & Wilson, M. A. Hippocampal replay of extended experience. Neuron 63, 497–507 (2009).

6 Diba, K. & Buzsáki, G. Forward and reverse hippocampal place-cell sequences during ripples. Nature Neuroscience 10, 1241–1242 (2007).

7 Foster, D. J. Replay comes of age. Annual Review of Neuroscience 40, 581–602 (2017).

8 Schwartenbeck, P. et al. Generative replay for compositional visual understanding in the prefrontal-hippocampal circuit. Cell (in press).

9 Liu, Y., Dolan, R. J., Kurth-Nelson, Z. & Behrens, T. E. J. Human replay spontaneously reorganizes experience. Cell 178, 640–652.e614 (2019).

10 Gupta, A. S., van der Meer, M. A. A., Touretzky, D. S. & Redish, A. D. Hippocampal replay Is not a simple function of experience. Neuron 65, 695–705 (2010).

11 Buzsaki, G. Hippocampal sharp wave−ripple: A cognitive biomarker for episodic memory and planning. Hippocampus 25, 1073–1188 (2015).

12 Ambrose, R. E., Pfeiffer, B. E. & Foster, D. J. Reverse replay of hippocampal place cells is uniquely modulated by changing reward. Neuron 91, 1124–1136 (2016).

13 Hahamy, A., Dubossarsky, H. & Behrens, T. E. J. The human brain reactivates context-specific past information at event boundaries of naturalistic experiences. Nature Neuroscience 26, 1080–1089 (2023).

14 Liu, Y., Mattar Marcelo, G., Behrens Timothy, E. J., Daw Nathaniel, D. & Dolan Raymond, J. Experience replay is associated with efficient nonlocal learning. Science 372, eabf1357 (2021).

15 Widloski, J. & Foster, D. J. Flexible rerouting of hippocampal replay sequences around changing barriers in the absence of global place field remapping. Neuron 110, 1547–1558 (2022).

16 Kaefer, K., Nardin, M., Blahna, K. & Csicsvari, J. Replay of behavioral sequences in the medial prefrontal cortex during rule switching. Neuron 106, 154–165 (2020).

17 Ji, D. & Wilson, M. A. Coordinated memory replay in the visual cortex and hippocampus during sleep. Nature Neuroscience 10, 100–107 (2007).

18 O’Neill, J., Boccara, C. N., Stella, F., Schönenberger, P. & Csicsvari, J. Superficial layers of the medial entorhinal cortex replay independently of the hippocampus. Science 355, 184–188 (2017).

19 Rudoy, J. D., Voss, J. L., Westerberg, C. E. & Paller, K. A. Strengthening individual memories by reactivating them during sleep. Science 326, 1079–1079 (2009).

20 Shanahan, L. K., Gjorgieva, E., Paller, K. A., Kahnt, T. & Gottfried, J. A. Odor-evoked category reactivation in human ventromedial prefrontal cortex during sleep promotes memory consolidation. elife 7, e39681 (2018).

21 Tambini, A. & Davachi, L. Awake reactivation of prior experiences consolidates memories and biases cognition. Trends in cognitive sciences 23, 876–890 (2019).

22 Schuck, N., W. & Niv, Y. Sequential replay of nonspatial task states in the human hippocampus. Science 364, eaaw5181 (2019).

23 Liu, Y., Nour, M. M., Schuck, N. W., Behrens, T. E. J. & Dolan, R. J. Decoding cognition from spontaneous neural activity. Nature Reviews Neuroscience 23, 204–214 (2022).

24 Kurth-Nelson, Z., Economides, M., Dolan, Raymond J. & Dayan, P. Fast sequences of non-spatial state representations in humans. Neuron 91, 194–204 (2016).

25 Yu, Q., et al. Reduced reverse replay in anxious individuals impairs reward prediction. bioRxiv (2023).

26 Wittkuhn, L. & Schuck, N. W. Dynamics of fMRI patterns reflect sub-second activation sequences and reveal replay in human visual cortex. Nature Communications 12, 1795 (2021).

27 Wittkuhn, L., Krippner, L. M. & Schuck, N. W. Statistical learning of successor representations is related to on-task replay. bioRxiv (2022).

28 Wimmer, G. E., Liu, Y., McNamee, D. C. & Dolan, R. J. Distinct replay signatures for prospective decision-making and memory preservation. Proceedings of the National Academy of Sciences 120, e2205211120 (2023).

29 Wimmer, G. E., Liu, Y., Vehar, N., Behrens, T. E. J. & Dolan, R. J. Episodic memory retrieval success is associated with rapid replay of episode content. Nature Neuroscience 23, 1025–1033 (2020).

30 Higgins, C. et al. Replay bursts in humans coincide with activation of the default mode and parietal alpha networks. Neuron 109, 882–893.e887 (2021).

31 Whittington, J. C. R., McCaffary, D., Bakermans, J. J. W. & Behrens, T. E. J. How to build a cognitive map: insights from models of the hippocampal formation. arXiv (2022).

32 Behrens, T. E. J. et al. What is a cognitive map? Organizing knowledge for flexible behavior. Neuron 100, 490–509 (2018).

33 Raichle, M. E. & Snyder, A. Z. A default mode of brain function: A brief history of an evolving idea. NeuroImage 37, 1083–1090 (2007).

34 Yeshurun, Y., Nguyen, M. & Hasson, U. The default mode network: where the idiosyncratic self meets the shared social world. Nature Reviews Neuroscience 22, 181–192 (2021).

35 Hassabis, D., Kumaran, D., Vann, S. D. & Maguire, E. A. Patients with hippocampal amnesia cannot imagine new experiences. Proceedings of the National Academy of Sciences 104, 1726–1731 (2007).

36 Constantinescu, A. O., O’Reilly, J. X. & Behrens, T. E. J. Organizing conceptual knowledge in humans with a gridlike code. Science 352, 1464–1468 (2016).

37 Park, S. A., Miller, D. S. & Boorman, E. D. Inferences on a multidimensional social hierarchy use a grid-like code. Nature Neuroscience 24, 1292–1301 (2021).

38 Baldassano, C., Hasson, U. & Norman, K. A. Representation of real-world event schemas during narrative perception. Journal of Neuroscience 38, 9689–9699 (2018).

39 Philiastides, M. G., Tu, T. & Sajda, P. Inferring macroscale brain dynamics via fusion of simultaneous EEG-fMRI. Annual Review of Neuroscience 44, 315–334 (2021).

40 Pisauro, M. A., Fouragnan, E., Retzler, C. & Philiastides, M. G. Neural correlates of evidence accumulation during value-based decisions revealed via simultaneous EEG-fMRI. Nature Communications 8, 15808 (2017).

41 Hauser, T. U. et al. The feedback-related negativity (FRN) revisited: New insights into the localization, meaning and network organization. NeuroImage 84, 159–168 (2014).

42 Liu, Y. et al. Temporally delayed linear modelling (TDLM) measures replay in both animals and humans. Elife 10, e66917 (2021).

43 Friston, K. J. et al. Psychophysiological and modulatory interactions in neuroimaging. NeuroImage 6, 218–229 (1997).

44 Nour, M. M., Liu, Y., Arumuham, A., Kurth-Nelson, Z. & Dolan, R. J. Impaired neural replay of inferred relationships in schizophrenia. Cell 184, 4315–4328.e4317 (2021).

45 McFadyen, J., Liu, Y. & Dolan, R. J. Differential replay of reward and punishment paths predicts approach and avoidance. Nature Neuroscience 26, 627–637 (2023).

46 Garvert, M. M., Dolan, R. J. & Behrens, T. E. J. A map of abstract relational knowledge in the human hippocampal–entorhinal cortex. eLife 6, e17086 (2017).

47 Karlsson, M. P. & Frank, L. M. Awake replay of remote experiences in the hippocampus. Nature Neuroscience 12, 913–918 (2009).

48 Ólafsdóttir, H. F., Bush, D. & Barry, C. The role of hippocampal replay in memory and planning. Curr Biol 28, R37–R50 (2018).

49 Klein-Flügge, M. C., Bongioanni, A. & Rushworth, M. F. S. Medial and orbital frontal cortex in decision-making and flexible behavior. Neuron 110, 2743–2770 (2022).

50 Schapiro, A. C., Turk-Browne, N. B., Norman, K. A. & Botvinick, M. M. Statistical learning of temporal community structure in the hippocampus. Hippocampus 26, 3–8 (2016).

51 Sherrill, K. R. et al. Generalization of cognitive maps across space and time. Cerebral Cortex 33, 7971–7992 (2023).

52 Silston, B. et al. Neural encoding of perceived patch value during competitive and hazardous virtual foraging. Nature Communications 12, 5478 (2021).

53 Baram, A. B., Muller, T. H., Nili, H., Garvert, M. M. & Behrens, T. E. J. Entorhinal and ventromedial prefrontal cortices abstract and generalize the structure of reinforcement learning problems. Neuron 109, 713–723.e717 (2021).

54 Vaidehi, S. N. et al. Stimulation of the posterior cingulate cortex impairs episodic memory encoding. The Journal of Neuroscience 39, 7173 (2019).

55 Bone, M. B. & Buchsbaum, B. R. Detailed episodic memory depends on concurrent reactivation of basic visual features within the posterior hippocampus and early visual cortex. Cerebral Cortex Communications 2, tgab045 (2021).

56 Favila, S. E., Lee, H. & Kuhl, B. A. Transforming the concept of memory reactivation. Trends in Neurosciences 43, 939–950 (2020).

57 Schapiro, A. C., McDevitt, E. A., Rogers, T. T., Mednick, S. C. & Norman, K. A. Human hippocampal replay during rest prioritizes weakly learned information and predicts memory performance. Nature Communications 9, 3920 (2018).

58 Agrawal, M., Mattar, M. G., Cohen, J. D. & Daw, N. D. The temporal dynamics of opportunity costs: A normative account of cognitive fatigue and boredom. Psychological review 129, 564 (2022).

59 Mattar, M. G. & Daw, N. D. Prioritized memory access explains planning and hippocampal replay. Nature Neuroscience 21, 1609–1617 (2018).

60 Dijkstra, N. & Fleming, S. M. Subjective signal strength distinguishes reality from imagination. Nature Communications 14, 1627 (2023).

61 Kaplan, R. et al. Hippocampal sharp-wave ripples influence selective activation of the default mode network. Curr Biol 26, 686–691 (2016).

62 Aronov, D., Nevers, R. & Tank, D. W. Mapping of a non-spatial dimension by the hippocampal–entorhinal circuit. Nature 543, 719–722 (2017).

63 Hafting, T., Fyhn, M., Molden, S., Moser, M.-B. & Moser, E. I. Microstructure of a spatial map in the entorhinal cortex. Nature 436, 801–806 (2005).

64 Ritvo, V. J. H., Turk-Browne, N. B. & Norman, K. A. Nonmonotonic plasticity: how memory retrieval drives learning. Trends in cognitive sciences 23, 726–742 (2019).

65 Peirce, J. et al. PsychoPy2: Experiments in behavior made easy. Behavior research methods 51, 195–203 (2019).

66 Allen, P. J., Josephs, O. & Turner, R. A method for removing imaging artifact from continuous EEG recorded during functional MRI. Neuroimage 12, 230–239 (2000).

67 Delorme, A. & Makeig, S. EEGLAB: an open source toolbox for analysis of single-trial EEG dynamics including independent component analysis. Journal of Neuroscience Methods 134, 9–21 (2004).

68 Esteban, O. et al. fMRIPrep: a robust preprocessing pipeline for functional MRI. Nature Methods 16, 111–116 (2019).

69 Gorgolewski, K. et al. Nipype: a flexible, lightweight and extensible neuroimaging data processing framework in python. Frontiers in Neuroinformatics 5, 13 (2011).

70 Abraham, A. et al. Machine learning for neuroimaging with scikit-learn. Frontiers in Neuroinformatics 8, 14 (2014).

71 Tustison, N. J. et al. N4ITK: improved N3 bias correction. IEEE Transactions on Medical Imaging 29, 1310–1320 (2010).

72 Avants, B. B., Epstein, C. L., Grossman, M. & Gee, J. C. Symmetric diffeomorphic image registration with cross-correlation: evaluating automated labeling of elderly and neurodegenerative brain. Medical Image Analysis 12, 26–41 (2008).

73 Zhang, Y., Brady, M. & Smith, S. Segmentation of brain MR images through a hidden Markov random field model and the expectation-maximization algorithm. IEEE Transactions on Medical Imaging 20, 45–57 (2001).

74 Dale, A. M., Fischl, B. & Sereno, M. I. Cortical surface-based analysis: I. Segmentation and surface reconstruction. Neuroimage 9, 179–194 (1999).

75 Klein, A. et al. Mindboggling morphometry of human brains. PLoS Computational Biology 13, e1005350 (2017).

76 Fonov, V. S., Evans, A. C., McKinstry, R. C., Almli, C. R. & Collins, D. L. Unbiased nonlinear average age-appropriate brain templates from birth to adulthood. NeuroImage 47, S102 (2009).

77 Jenkinson, M., Bannister, P., Brady, M. & Smith, S. Improved optimization for the robust and accurate linear registration and motion correction of brain images. Neuroimage 17, 825–841 (2002).

78 Cox, R. W. & Hyde, J. S. Software tools for analysis and visualization of fMRI data. NMR in Biomedicine: An International Journal Devoted to the Development and Application of Magnetic Resonance In Vivo 10, 171–178 (1997).

79 Greve, D. N. & Fischl, B. Accurate and robust brain image alignment using boundary-based registration. Neuroimage 48, 63–72 (2009).

80 Power, J. D. et al. Methods to detect, characterize, and remove motion artifact in resting state fMRI. Neuroimage 84, 320–341 (2014).

81 Behzadi, Y., Restom, K., Liau, J. & Liu, T. T. A component based noise correction method (CompCor) for BOLD and perfusion based fMRI. Neuroimage 37, 90–101 (2007).

82 Satterthwaite, T. D. et al. An improved framework for confound regression and filtering for control of motion artifact in the preprocessing of resting-state functional connectivity data. Neuroimage 64, 240–256 (2013).

83 Lanczos, C. Evaluation of noisy data. Journal of the Society for Industrial and Applied Mathematics, Series B: Numerical Analysis 1, 76–85 (1964).

84 Pedregosa, F. et al. Scikit-learn: Machine learning in Python. Journal of Machine Learning Research 12, 2825–2830 (2011).

85 Smith, S. M. et al. Advances in functional and structural MR image analysis and implementation as FSL. Neuroimage 23, S208–S219 (2004).

86 Poldrack, R. A. Region of interest analysis for fMRI. Social Cognitive and Affective Neuroscience 2, 67–70 (2007).

87 Fischl, B. et al. Automatically parcellating the human cerebral cortex. Cerebral Cortex 14, 11–22 (2004).

88 Haxby, J. V. et al. Distributed and overlapping representations of faces and objects in ventral temporal cortex. Science 293, 2425–2430 (2001).

89 Kunz, L., Deuker, L., Zhang, H. & Axmacher, N. in Handbook of Behavioral Neuroscience Vol. 28 Handbook of in Vivo Neural Plasticity Techniques In (ed Denise Manahan-Vaughan) 481–508 (Elsevier, 2018).

90 Desikan, R. S. et al. An automated labeling system for subdividing the human cerebral cortex on MRI scans into gyral based regions of interest. NeuroImage 31, 968–980 (2006).

91 Eickhoff, S. B. et al. A new SPM toolbox for combining probabilistic cytoarchitectonic maps and functional imaging data. NeuroImage 25, 1325–1335 (2005).

92 Benjamini, Y. Discovering the false discovery rate. Journal of the Royal Statistical Society: Series B (Statistical Methodology) 72, 405–416 (2010).

93 Lifanov, J., et al. Reconstructing spatio-temporal trajectories of visual object memories in the human brain. bioRxiv (2022).

